# The *Toxoplasma* subpellicular network is highly interconnected and defines parasite shape for efficient motility and replication

**DOI:** 10.1101/2023.08.10.552545

**Authors:** Peter S. Back, Vignesh Senthilkumar, Charles P. Choi, Andrew M. Ly, Anne K. Snyder, Justin G. Lau, Gary E. Ward, Peter J. Bradley

**Affiliations:** Molecular Biology Institute, University of California, Los Angeles, United States of America; Department of Microbiology, Immunology, and Molecular Genetics, University of California, Los Angeles, United States of California; Department of Microbiology and Molecular Genetics, Larner College of Medicine, University of Vermont, Burlington, Vermont, United States of America

**Keywords:** Alveolata, Apicomplexa, alveolin, subpellicular network, photoreactive crosslinking

## Abstract

Apicomplexan parasites possess several specialized structures to invade their host cells and replicate successfully. One of these is the inner membrane complex (IMC), a peripheral membrane-cytoskeletal system underneath the plasma membrane. It is composed of a series of flattened, membrane-bound vesicles and a cytoskeletal subpellicular network (SPN) comprised of intermediate filament-like proteins called alveolins. While the alveolin proteins are conserved throughout the Apicomplexa and the broader Alveolata, their precise functions and interactions remain poorly understood. Here, we describe the function of one of these alveolin proteins, TgIMC6. Disruption of IMC6 resulted in striking morphological defects that led to aberrant motility, invasion, and replication. Deletion analyses revealed that the alveolin domain alone is largely sufficient to restore localization and partially sufficient for function. As this highlights the importance of the IMC6 alveolin domain, we implemented unnatural amino acid photoreactive crosslinking to the alveolin domain and identified multiple binding interfaces between IMC6 and two other cytoskeletal proteins – IMC3 and ILP1. To our knowledge, this provides the first direct evidence of protein-protein interactions in the alveolin domain and supports the long-held hypothesis that the alveolin domain is responsible for filament formation. Collectively, our study features the conserved alveolin proteins as critical components that maintain the parasite’s structural integrity and highlights the alveolin domain as a key mediator of SPN architecture.

## Introduction

The superphylum Alveolata contains a remarkably diverse group of protozoans, including the free-living ciliates and dinoflagellates as well as the parasitic apicomplexans. Despite these lifestyle differences, the alveolates are unified by a peripheral system of membranes underlying the plasma membrane called the alveoli and a conserved group of proteins called the alveolins [1]. These two features define the superphylum and highlight the alveoli as critical cellular structures for the survival of these diverse organisms. Of these, the apicomplexans have garnered the most attention due to their severe public health and economic burdens [2]. Notable human pathogens include *Toxoplasma gondii* (toxoplasmosis), *Plasmodium* spp. (malaria), and *Cryptosporidium* spp. (cryptosporidiosis), which together cause an enormous disease burden globally that results in a tremendous number of fatalities [3–5]. Veterinary pathogens include *Neospora* spp. and *Eimeria* spp., which cause large numbers of disease in livestock and subsequent economic losses [6,7].

In the Apicomplexa, the alveoli are called the inner membrane complex (IMC) and is situated underneath the plasma membrane as in other alveolates. The IMC has been studied most extensively in *T. gondii* and *Plasmodium* spp. and contains three main functions. It first serves as a platform for the glideosome, an actin-myosin motor that powers gliding motility and host cell invasion [8]. Second, it provides a scaffold for daughter cell assembly throughout the replication process [9–11]. Finally, the apical cap region of the IMC houses the regulatory center for cytoskeletal disassembly during the final stages of replication [12–14]. While these functions are generally conserved in other apicomplexans, it is unlikely that they are conserved in other alveolates. Due to their non-parasitic lifestyle, ciliates and dinoflagellates have likely co-opted their respective alveoli to meet the demands of a free-living environment. Thus, determining the precise roles of the alveolin proteins promises to provide insights into these differences.

In apicomplexans, the alveolins are believed to be the major constituents of the subpellicular network (SPN) [15–21,13,12]. The SPN is a highly interwoven mesh of filaments that underlies the membrane vesicles of the IMC and provides a cytoskeletal foundation for the parasite [9,18]. The alveolins are categorized by the presence of a conserved alveolin domain, a region of the protein containing proline and valine-rich repeats [1]. It is speculated that the alveolin domain mediates the formation of filaments via protein-protein interactions that ultimately establish the SPN. However, direct experimental evidence is lacking, largely due to the detergent insoluble nature of these proteins that limit the use of standard interaction methods such as co-immunoprecipitation. To overcome this barrier, we recently adapted unnatural amino acid (UAA) photocrosslinking to *T. gondii*, which identifies the interacting partner within the native environment of the parasite [22]. This approach uses the zero-length crosslinker *p*-azidophenylalanine (Azi), enabling us to map specific binding interfaces that provide structural information regarding the interaction [23]. We previously used this technique to determine that TgILP1 binds two alveolin proteins, IMC3 and IMC6. However, these interactions were mapped to their variable N and C terminal regions rather than the core alveolin domains. Thus, the alveolin domains remain unexplored in this family of proteins. Functionally, only some of the alveolin proteins have been studied so far. Of these, many were shown to play a role in replication or in providing tensile strength, but none were shown to be important or essential for parasite fitness [24,25]. In contrast, the alveolins that have eluded characterization are typically those with low genome-wide CRISPR screen (GWCS) phenotype scores, suggesting essentiality [26].

In this study, we report the successful knockout of the alveolin TgIMC6. With this breakthrough, we elucidate its critical role in maintaining parasite shape and describe the downstream consequences of its absence on parasite motility and host cell invasion. We also characterize the severe replication errors exhibited by Δ*imc6* parasites. To determine the significance of the alveolin domain, we use deletion analyses to demonstrate that the alveolin domain is largely sufficient to restore IMC6 localization and function. We then implement the recently developed photoreactive crosslinking approach to pinpoint direct interactions and map the binding interfaces between the IMC6 alveolin domain and other cytoskeletal proteins of the SPN. This firmly establishes the alveolin domain as a major region of protein-protein contact, supporting the long-held hypothesis that the alveolin proteins mediate the formation of SPN filaments.

## Results

### Disrupting IMC6 causes severe defects *in vitro* and *in vivo*

Of the alveolin proteins with low phenotype scores, IMC6 was assigned the highest score of -3.19 [26,27]. Recent work from our lab demonstrated that genes with similar phenotype scores (ISAP1: -3.49 and IMC29: -3.95) could be disrupted [28,29]. Thus, we wanted to determine if IMC6 could also be knocked out. As previously reported, IMC6 localizes to the IMC of both maternal and daughter parasites with enrichment in the daughter buds (Fig 1A) [18]. We were indeed successful in generating a knockout strain (Δ*imc6*), verified by the absence of protein expression using immunofluorescence assay (IFA) and by recombination at the genomic locus using PCR (Fig 1B and 1C). We then generated a complementation construct with the full-length IMC6 cDNA driven by its endogenous promoter with a C-terminal 1xV5 epitope tag (Fig 1D). Expressing this construct in Δ*imc6* parasites restored protein localization and expression similar to wild-type levels as determined by IFA and western blot (complemented strain: IMC6c, Fig 1E and 1F).

**Figure 1.**
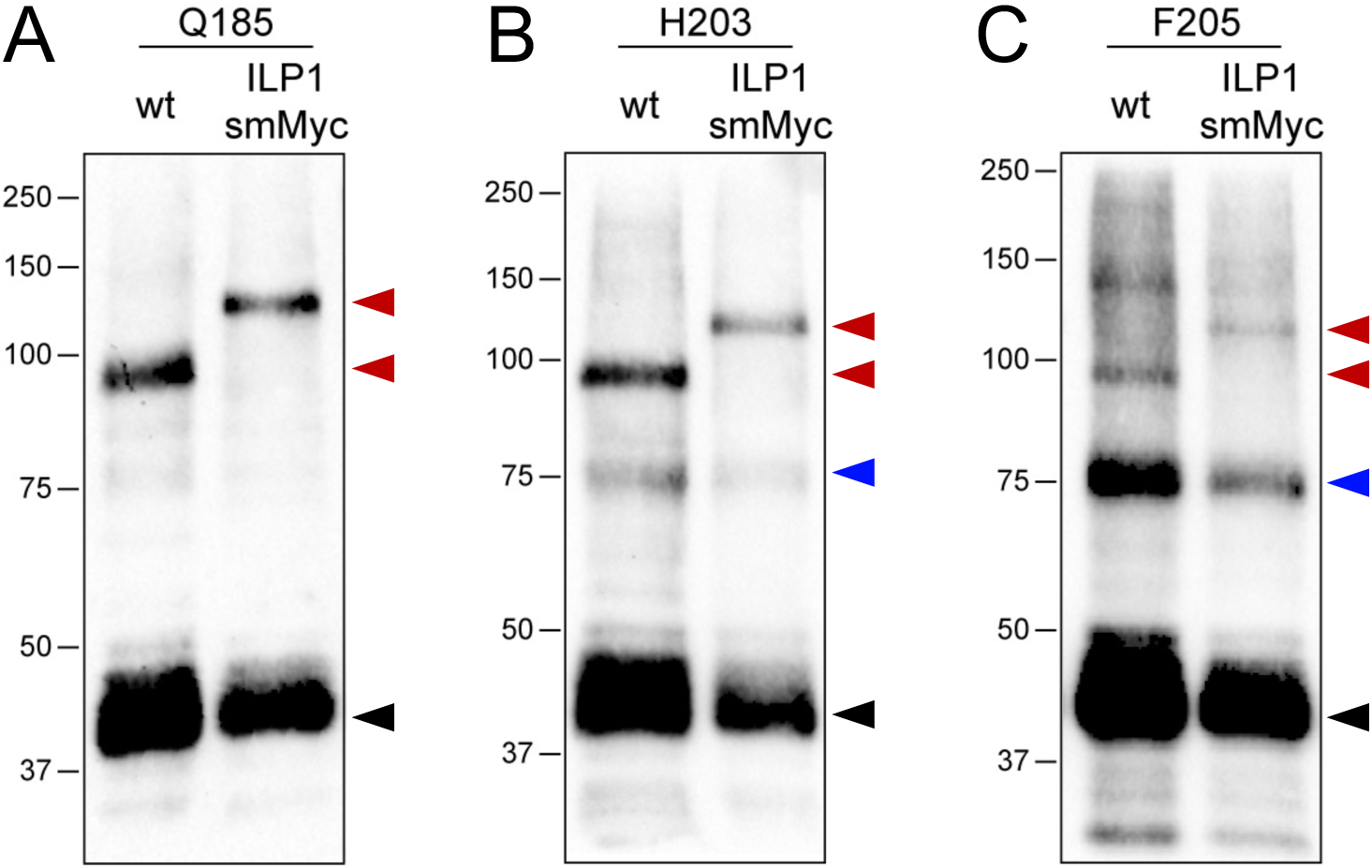
Disrupting IMC6 causes severe fitness and virulence defects. (A) Immunofluorescence assay (IFA) of wild-type parasites showing proper IMC localization in mature and budding parasites. Magenta, anti-IMC6; green, anti-IMC10. (B) IFA of Δ*imc6* parasites showing absence of IMC6 in mature and budding parasites. Magenta, anti-IMC6; green, anti-IMC10. (C) PCR verification of the genomic loci from WT (RHΔ*hxgprt*) and Δ*imc6* parasites. Diagram illustrates primers used to amplify the IMC6 coding sequence (blue arrows) and the site of recombination for the knockout locus (magenta arrows). (D) Diagram of the complementation construct, which includes the full-length IMC6, the endogenous promoter and a 1xV5 epitope tag. The alveolin domain is highlighted in blue. (E) IFA of IMC6c parasites showing restored localization of IMC6. Magenta, anti-V5; green, anti-IMC3. (F) Western blot of whole cell lysates illustrating the absence (Δ*imc6*) and rescue (IMC6c) of IMC6 expression. The difference in migration between WT and IMC6c is due to the 1xV5 epitope tag. IMC6 was detected with anti-IMC6; ROP13 was used as a loading control and detected with anti-ROP13. (G) Quantification of plaque areas depicting the severe growth defect of Δ*imc6* parasites, which is fully restored in IMC6c parasites. Triplicates performed with 30-40 plaques measured per replicate. Significance was determined using two-way ANOVA. ****: p < 0.0001. (H) Quantification of plaque efficiency as the percentage of plaques over the total number of parasites added to each replicate (triplicates performed). Significance was determined using multiple two-tailed t tests. ****: p < 0.0001. (I) Virulence assay using four mice highlighting the dramatic virulence defect of Δ*imc6* parasites (black line). All scale bars are 2 µm.

To assess the overall fitness cost of disrupting *IMC6*, we first performed plaque assays. Measuring both plaque area and plaque-forming efficiency indicated a severe 78.8% reduction in plaque size and a 71.7% reduction in plaque efficiency (Fig 1G and H). These defects were fully rescued in the IMC6c parasites. To determine if virulence is also affected, we infected mice with 10^2^ wild-type, 10^5^ Δ*imc6*, or 10^2^ IMC6c parasites (Fig 1I). As expected, the mice injected with wild-type or IMC6c parasites succumbed to the infection after 7 days. In contrast, of the four mice injected with a 1000-fold greater number of knockout parasites, three survived the infection. This demonstrates a dramatic decrease in virulence that corroborates the *in vitro* growth defects and highlights the importance of IMC6 for parasite fitness.

### Δ*imc6* parasites exhibit gross morphological defects

Upon disrupting IMC6, one of the most striking changes was parasite morphology. Δ*imc6* parasites appeared grossly misshapen in both intracellular vacuoles and extracellular parasites (Fig 2A and 2B). To quantify this shape defect in a nonbiased manner, we used ImageStream imaging flow cytometry [21]. We measured >20,000 individual extracellular parasites for each strain and evaluated parasite morphology under two main categories – aspect ratio and circularity (Fig 2C and D). As expected, the aspect ratios of wild-type parasites indicated an elongated morphology with a median of 0.59. In contrast, the aspect ratios of knockout parasites were higher with a median of 0.78, indicating a substantially more rounded shape on average. This shape defect was fully restored in the complemented parasites. Similarly for circularity, the wild-type and IMC6c populations contained, on average, less circular cells with medians of 4.7 and 5.1, respectively. In contrast, the circularity values for Δ*imc6* parasites indicated substantially rounder cells with a median of 7.8, again highlighting the grossly misshapen morphology of these knockout parasites.

**Figure 2.**
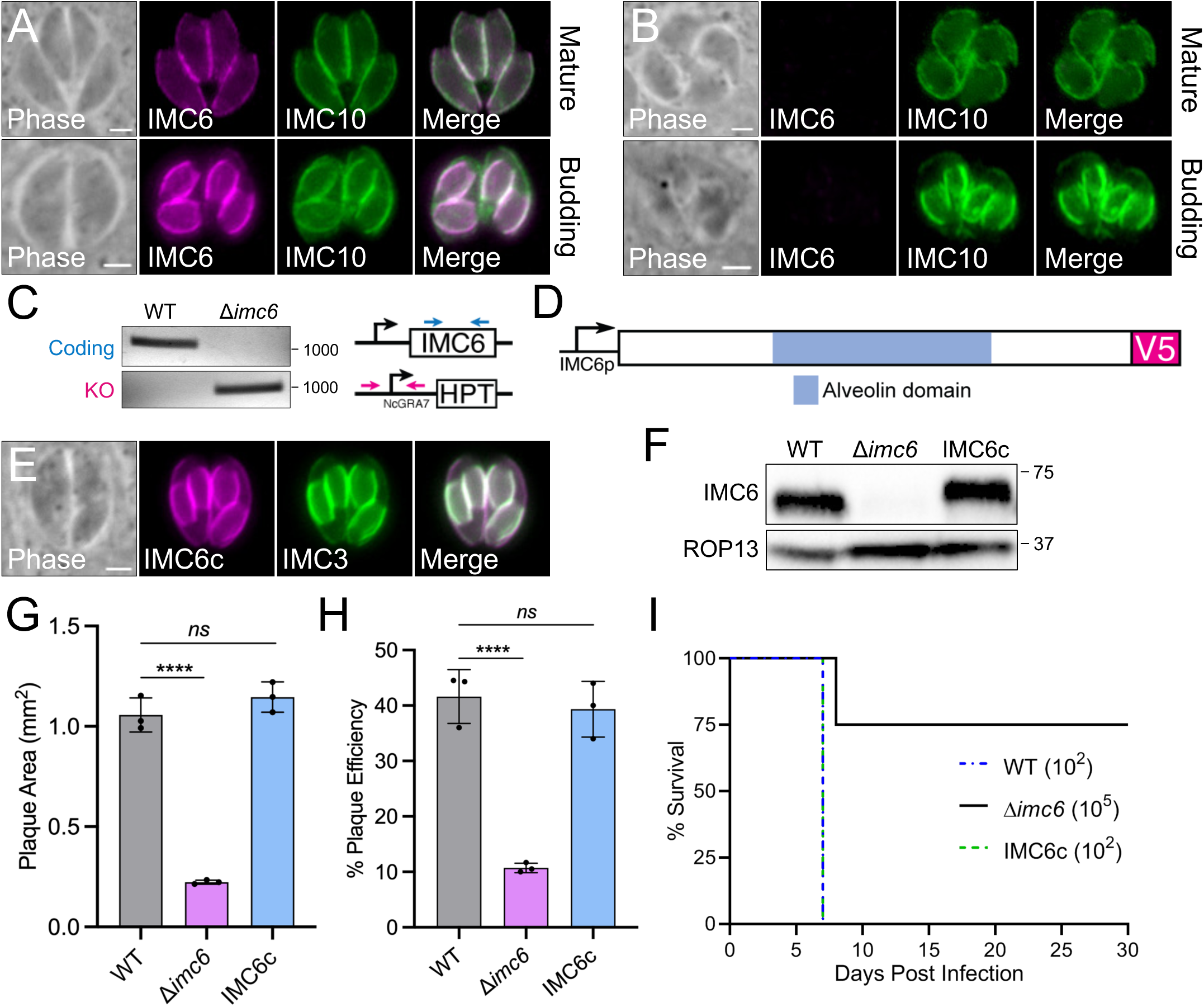
Δ*imc6* parasites are extremely misshapen. (A) IFA of a representative field of Δ*imc6* vacuoles, illustrating the grossly swollen parasites. Magenta, anti-ISP1; green, anti-IMC3. (B) Representative phase contrast images of a field of extracellular WT and Δ*imc6* parasites, highlighting the extremely rounded shape of Δ*imc6* parasites. (C) ImageStream flow cytometry analysis showing the aspect ratios of extracellular WT, Δ*imc6* and IMC6c parasite populations. Greater aspect ratios indicate rounder objects. Median aspect ratio for WT is 0.59, Δ*imc6* is 0.78, and IMC6c is 0.61. (D) Imagestream analysis using the circularity parameter, where greater values indicate rounder objects. Median circularity value for WT is 4.72, Δ*imc6* is 7.78, and IMC6c is 5.05. (E-F) Bivariate plots of the Circularity feature vs. the Elongated Machine Learning (ML) classifier on the WT (E) and Δ*imc6* (F) samples (see S1 Fig). The WT population contains 85.3% elongated and 13.3% circular objects. The Δ*imc6* population contains 46.6% elongated and 46.5% circular objects. This population shift is highlighted by the density plots. The assigned values indicate confidence given by the Elongated ML classifier (higher values means greater confidence of elongatedness).

To extend this quantification, we supplemented the circularity measurement with the Elongated Machine Learning (EloML) classifier. The EloML classifier considers a set of 10 features and assigns a binary classification for each object as either elongated or circular (Fig S1). Taught on 200 wild-type and 200 knockout parasites, the EloML classifier was designed to create a threshold for elongated/circular objects and to assign a confidence value for each one. Upon analyzing our parasites with this approach, the wild-type strain contained 85.3% elongated and 13.3% circular populations (Fig 2E). In contrast, the Δ*imc6* strain contained 46.6% elongated and 46.5% circular populations (Fig 2F). This dramatic shift in the population is illustrated by the density scatter plots. Taken together, imaging flow cytometry permits a robust analysis of morphological defects and demonstrates that IMC6 is critical to maintain the parasite’s overall shape and structural integrity.

### Shape-derived motility defects result in delayed parasite invasion

As parasite morphology has been shown to affect gliding motility, we evaluated Δ*imc6* parasites using 3D motility assays [30]. Fig 3A shows representative maximum intensity projections (MIPs) of parasite motility in the Matrigel-based environment, illustrating the proportion of motile parasites, track length, and track shape. To quantify these parameters, motility images were captured over an 80-second period. Using a 2.8-µm minimum displacement threshold (see methods), we observed no significant difference in the proportion of parasites moving or track length between wild-type and knockout parasites (Fig 3B and C). However, the Δ*imc6* parasites exhibited a modest but significant defect in maximum speed achieved along their trajectories compared to wild-type parasites (Fig 3D). We also noted differences in trajectory shape. For wild-type parasites, the representative MIPs and videos depict the characteristic asymmetrical helical shape of the trajectory (Fig 3A, S1 Video, and S2 Video) [30]. Many of the Δ*imc6* trajectories, however, show reduced helicity, ranging from no apparent helicity to a helix with a much longer period (*i.e.*, a “flatter” corkscrew shape). We were unable to quantify these differences in shape due to the inconsistencies in the Δ*imc6* tracks (Fig. 3A, S3-S6 Video). Nevertheless, both the decrease in maximum trajectory speed and apparent differences in trajectory shape were reversed in the complemented strain (Fig. 3A-D, S7 Video, and S8 Video). Together, these assays indicate that motility is significantly affected in the Δ*imc6* parasites. To assess whether these motility defects are due to problems in microneme secretion, we performed a microneme secretion assay and found that exocytosis remains unperturbed in the Δ*imc6* parasites (S2 Fig). This suggests that the motility defects are primarily caused by aberrant parasite shape, highlighting the biophysical requirements of gliding motility.

**Figure 3.**
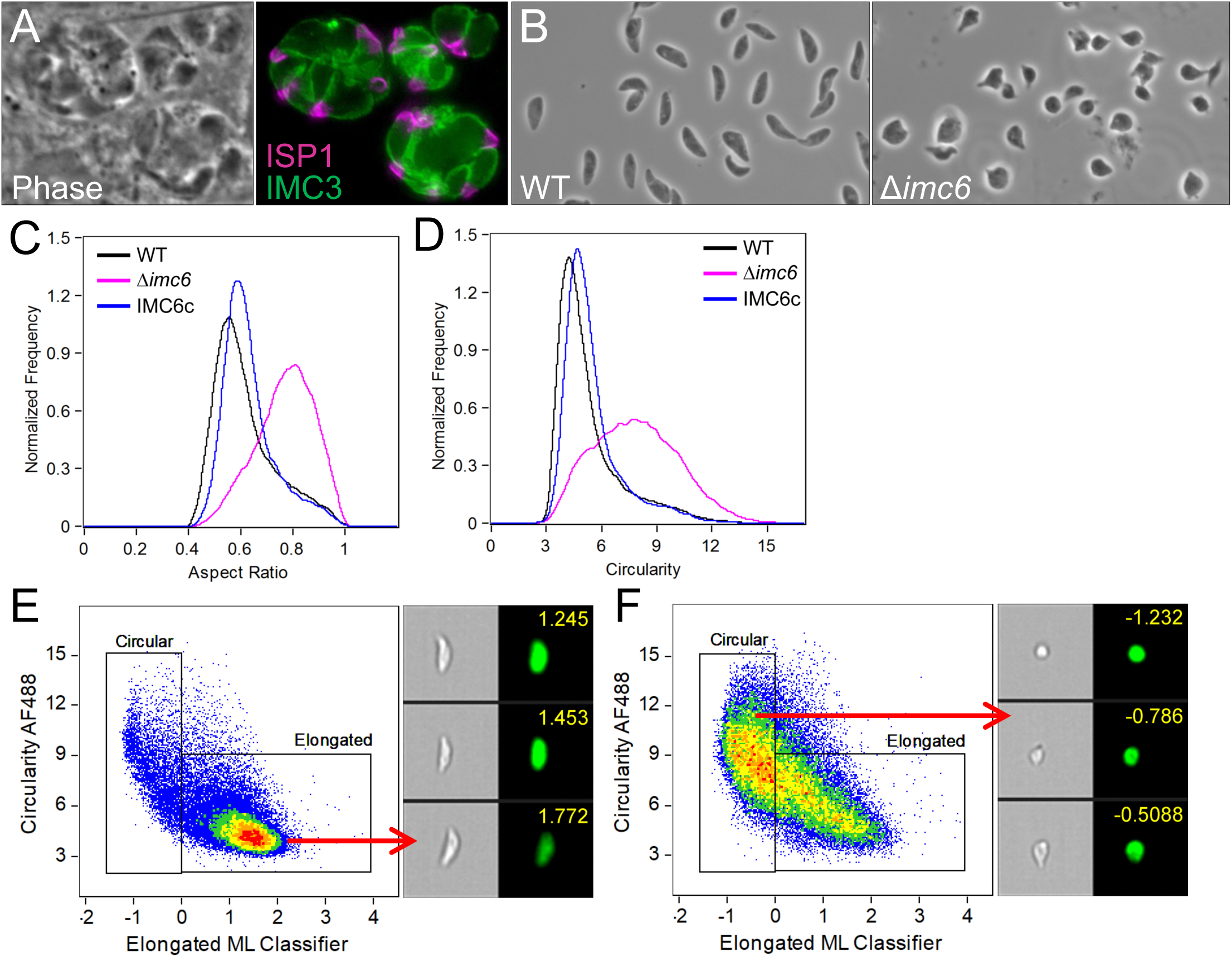
Δ*imc6* parasites exhibit slower motility and delayed invasion. (A) Representative maximum intensity projections showing trajectories of WT, Δ*imc6*, and IMC6c parasites in Matrigel, imaged over 80 seconds. Greyscale images were inverted to better visualize parasite trajectories. N indicates the number of tracks quantified for this replicate. Scale bars are 25 µm. (B-D) Quantifications of percent moving, track length, and maximum track speed. All trajectory data were acquired from three biological replicates, each consisting of three technical replicates. Significance was determined using one-way ANOVA. *: p = 0.0457. (E-G) Red-green invasion assays with 5-min (E), 20-min (F) and 60-min (G) invasion permissive conditions. Magenta represents attached and green represents invaded parasites. Note the increasing trend for the number of invaded Δ*imc6* parasites as invasion times increase. Triplicate experiments were performed by counting the number of parasites per host nucleus (25 host nuclei across at least 7 different fields per replicate). Significance was calculated using two-way ANOVA. ****: p < 0.0001.

As motility is intimately linked to invasion, we wanted to determine if decreased motility speed impacts invasion efficiency. To test this, we adapted the standard invasion assays by modulating the time spent in invasion-permissive conditions. With a 5-minute invasion period, the number of Δ*imc6* parasites invaded (1.3±0.3 parasites/host cell) was significantly fewer compared to that of wild-type parasites (4.2±0.6 parasites/host cell) while the total number of parasites attached and invaded remained comparable (Fig 3E). Thus, the ability to attach to a host cell appears unaffected but the efficiency of invasion is severely hampered. We hypothesized that slower speeds observed in the 3D motility assay may be compensated by increased invasion time. To test this, we performed invasion assays with 20-and 60-minute invasion periods (Fig 3F and 3G). Consistent with our hypothesis, the number of invaded Δ*imc6* parasites increased significantly at the 20-minute and even more at the 60-minute time points (3.7±0.1 and 5.8±0.8 parasites/host cell, respectively). This indicates that the knockout parasites can largely catch up to the invasion efficiency of wild-type parasites given enough time, firmly linking parasite shape to motility and invasion.

### IMC6 is important for proper parasite replication

As the subtle defects in motility and invasion are likely insufficient to produce the severe plaque defects of Δ*imc6* parasites, we evaluated whether replication is also affected since the localization of IMC6 is enriched in daughter buds. We first assessed overall replication by quantifying the number of parasites per vacuole at 24 and 32 hours post infection (hpi) (Fig 4A and 4B). At 24 hpi, the majority of wild-type vacuoles contained 8 parasites. In contrast, Δ*imc6* vacuoles were more evenly split between 4 and 8 parasites per vacuole, indicating they are progressing significantly slower compared to wild-type parasites. This defect became increasingly pronounced at 32 hpi, where the majority of wild-type vacuoles contained 16 parasites while a significantly fewer number of Δ*imc6* vacuoles contained 16 (Fig 4B). Thus, Δ*imc6* parasites can progress through endodyogeny, albeit at a considerably slower rate than wild-type parasites.

**Figure 4.**
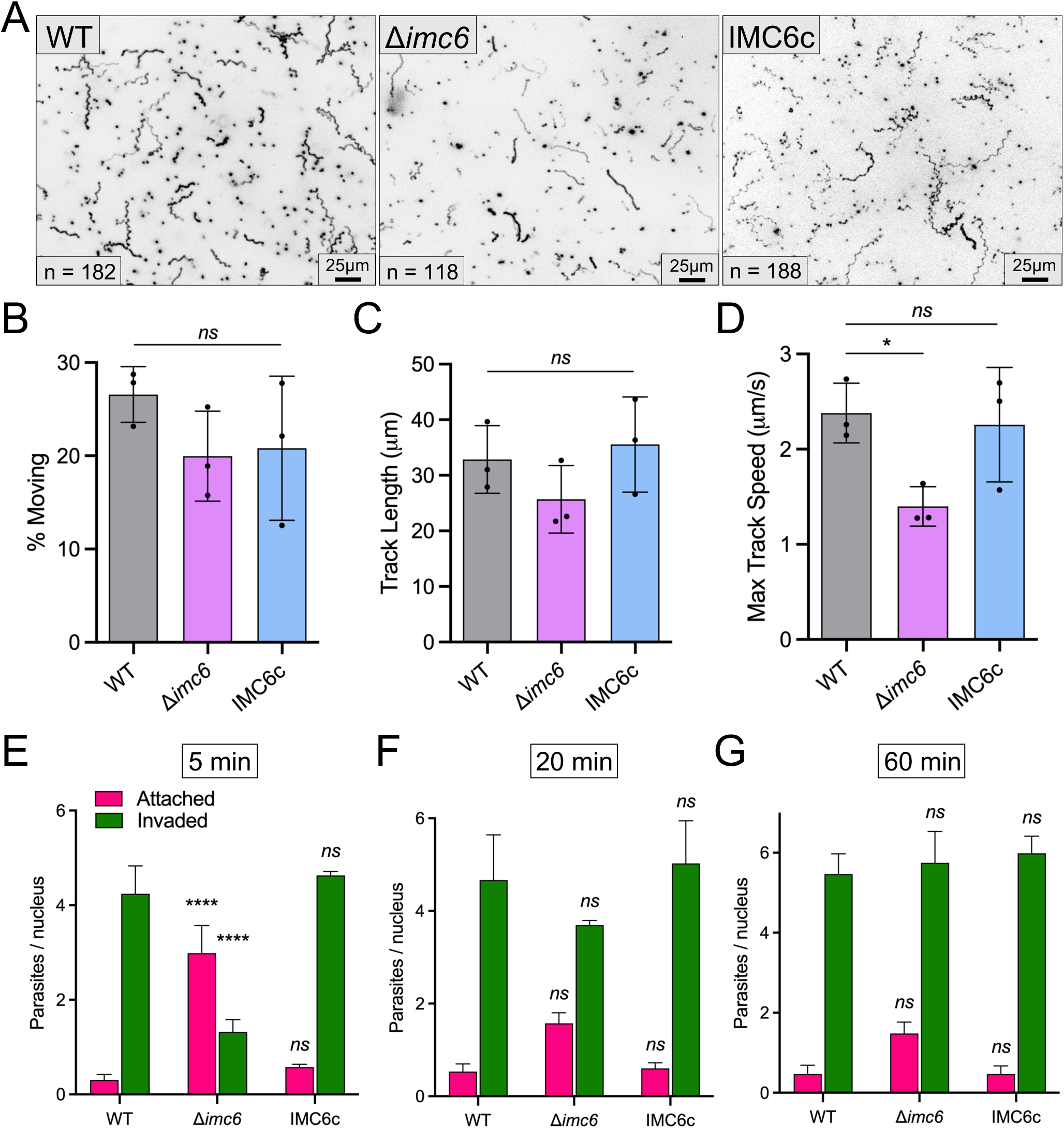
IMC6 is important for faithful endodyogeny. (A-B) Replication assays conducted at 24-hours (A) and 32-hours (B) post infection. Triplicate experiments were performed by quantifying the number of parasites/vacuole with >300 total vacuoles counted across at least 15 different fields per replicate. Significance was calculated using two-way ANOVA. (A) includes significance for 4 and 8 parasites per vacuole. (B) includes significance for 4, 8, and 16 parasites per vacuole. **: p = 0.0018. ****: p < 0.0001. (C-G) Representative IFAs depicting the various replication defects as indicated on the right side of each panel. Magenta, anti-ISP1; green, anti-IMC3. Scale bars are 2 µm. (H) Quantification of abnormal vacuoles as categorized by the presence of any one of the replication defects. Triplicate experiments were performed with >200 total vacuoles counted across at least 15 different fields per replicate. Significance was calculated using two-tailed t tests. ****: p < 0.0001. (I) Representative field of extracellular Δ*imc6* parasites highlighting the presence of incompletely separated parasites (white arrows). Magenta, anti-ISP1; green, anti-IMC3. Scale bar is 5 µm. (J) Quantification of the incomplete separation phenotype in extracellular parasites. Triplicates performed by categorizing >300 individual parasites across at least 15 different fields per replicate. Significance was calculated using two-tailed t tests. ****: p < 0.0001.

To pinpoint specific replication defects, we first stained Δ*imc6* parasites with a series of markers to evaluate various parasite organelles. These IFAs indicated that the apicoplast, mitochondria, PLV/VAC, micronemes and rhoptries localize properly and appear unaffected in the absence of IMC6 (S3 Fig). We then stained parasites with the IMC markers ISP1 and IMC3 to assess the fidelity of endodyogeny. We noticed several errors including asynchronous division, >2 daughter buds per maternal parasite, grossly unorganized vacuoles, large breaks in the IMC structure, and incomplete separation (Fig 4C-4G). Many of the Δ*imc6* vacuoles exhibited multiple replication defects simultaneously. Thus, rather than quantifying each defect individually, we categorized each vacuole as either standard or abnormal depending on the presence of any one of these replication defects. This revealed an astonishing 92.8±6.1% of abnormal Δ*imc6* vacuoles compared to 7.4±2.7% of abnormal wild-type vacuoles and 4.7±1.4% of abnormal IMC6c vacuoles (Fig 4H). We were particularly intrigued by the incompletely separated parasites as this phenotype appears to be distinct from previously reported IMC mutants. In the final stages of endodyogeny, the parasites seem to stall very early during cytokinesis, resulting in conjoined bodies (Fig 4G). We even observed conjoined parasites that have begun the next round of division, suggesting that cytokinesis is decoupled from the cell cycle in these parasites (Fig 4G). This defect was also seen in naturally egressed extracellular parasites, with 19.4±3.5% of Δ*imc6* parasites exhibiting this phenotype (Fig 4I, J). Taken together, the cumulative effect of aberrant replication and less efficient invasion likely explains the severe growth defects *in vitro* and virulence defects *in vivo*.

### The alveolin domain is largely sufficient for IMC6 localization and function

We and others have previously reported that the alveolin domain is largely sufficient for the proper localization of IMC3, IMC6, and IMC8 [18,22]. However, these studies were done in wild-type parasites, so the functional significance of the alveolin domain has not been determined. Thus, we generated IMC6 deletion constructs that truncate the N-terminal and/or C-terminal regions flanking the alveolin domain and expressed them in Δ*imc6* parasites. We first assessed the localization of each truncated protein and found results consistent with our previous study. The localization of IMC6^2-290^ closely resembles that of the full-length IMC6 while both IMC6^128-290^ and IMC6^128-444^ exhibit slightly more cytoplasmic staining (Fig 5A-C). This demonstrates that the alveolin domain alone is sufficient to restore much of the protein’s localization.

**Figure 5.**
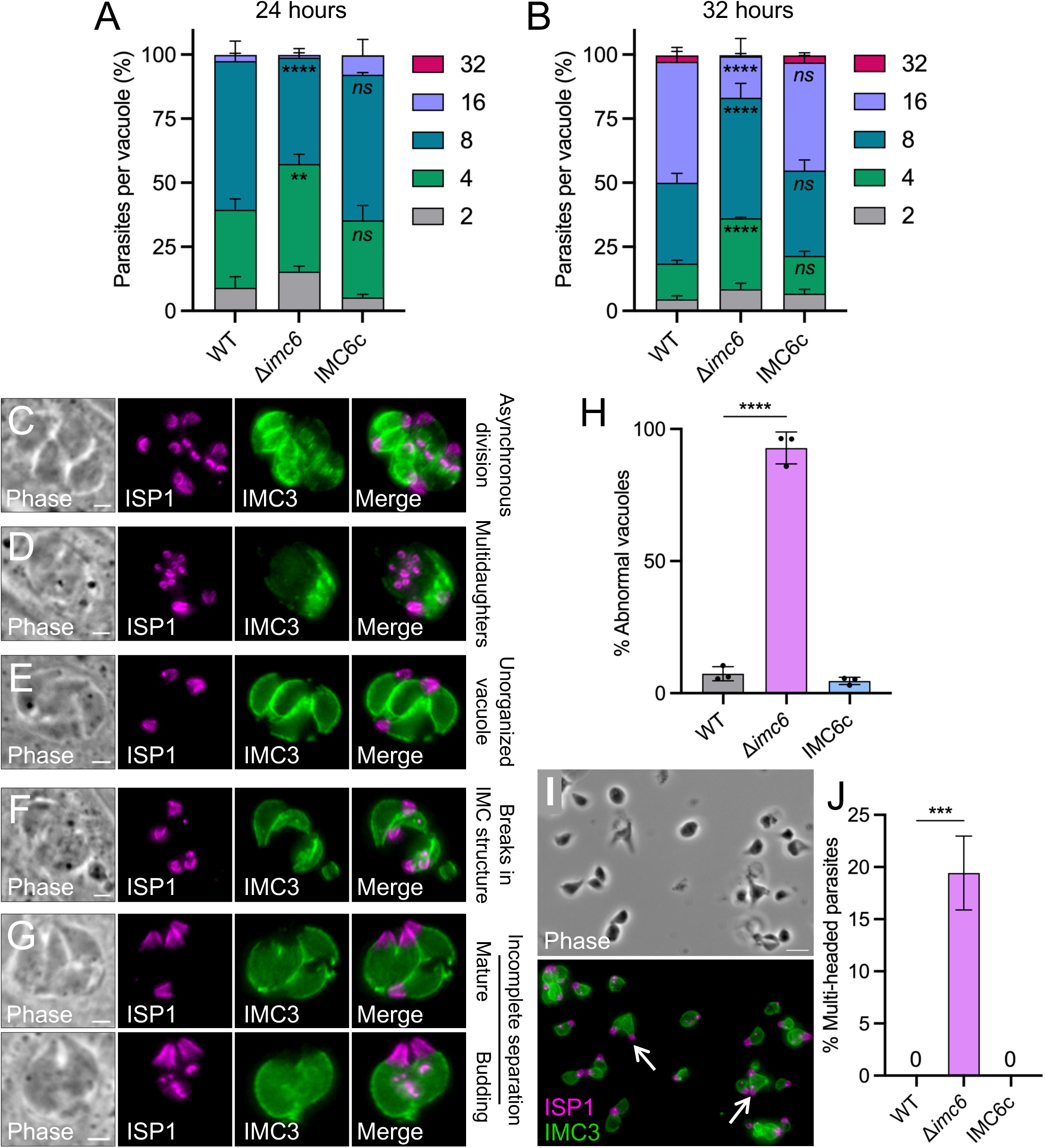
Domain analysis of IMC6 correlates parasite form and function. (A) Diagram and IFA of IMC6^2-290^, which truncates the region N-terminal to the alveolin domain, showing proper localization of IMC6^2-290^. Magenta, anti-V5; green, anti-IMC3. (B) Diagram and IFA of IMC6^128-290^, which truncates both the N-terminal and C-terminal regions of the protein. IMC6^2-290^ partially localizes to the IMC but shows additional cytoplasmic staining. Magenta, anti-V5; green, anti-IMC3. (C) Diagram and IFA of IMC6^128-444^, which truncates the region C-terminal regions to the alveolin domain, shows partial localization to the IMC with additional cytoplasmic staining. Magenta, anti-V5; green, anti-IMC3. (D) Quantification of plaque assay areas highlighting the partial rescue of growth defect by the three different deletion constructs. The data for WT, Δ*imc6*, and IMC6c were collected in the same experiment as those from Fig 1G and are shown again here to facilitate a direct comparison. Triplicates were performed by measuring 30-40 plaques per replicate, and significance was calculated using two-way ANOVA. ****: p < 0.0001. (E) Quantification of plaque efficiency showing partial rescue by each deletion construct. Note the increasing trend for IMC6^2-290^. The data for WT, Δ*imc6*, and IMC6c were collected in the same experiment as those from Fig 1H and are shown again here to facilitate a direct comparison. Significance was calculated using two-tailed t tests. ****: p < 0.0001. (F-G) ImageStream analysis of parasite populations expressing the three different deletion constructs, with (F) showing aspect ratio and (G) showing circularity parameters. The data for WT, Δ*imc6*, and IMC6c were collected in the same experiment as those from Fig 2C and 2D and are shown again here for ease of comparison. Median aspect ratio for IMC6^2-290^ is 0.68, IMC6^128-290^ is 0.73, and IMC6^128-444^ is 0.73. Median circularity value for IMC6^2-290^ is 6.04, IMC6^128-290^ is 6.63, and IMC6^128-444^ is 6.71. All scale bars are 2 µm.

To dissect the function of each domain, we performed plaque assays. Measuring both plaque area and plaque efficiency revealed a pattern that mimics the localization data, in which IMC6^2-290^ almost completely rescues the plaque defect while IMC6^128-290^ and IMC6^128-444^ only partially rescues the plaque defect (Fig 5D and 5E). We then used ImageStream to determine if localization and function are linked to parasite shape. Consistent with the plaque assays, the quantifications for aspect ratio and circularity followed a similar pattern (Fig 5F and 5G). Parasites expressing IMC6^2-290^ most resembled the elongated shape of wild-type parasites, though not fully. In contrast, parasites expressing IMC6^128-290^ or IMC6^128-444^ were moderately rounded, indicating a partial rescue of shape. Taken together, this domain analysis demonstrates that the alveolin domain alone is largely sufficient for localization and function with the N-terminal third of the protein contributing a partial role. This highlights the significance of the alveolin domain, and importantly, uncovers a strong correlation between parasite shape and parasite fitness, with more rounded parasites exhibiting decreased fitness. Thus, IMC6 provides a critical structural role that defines parasite shape, which ultimately impacts parasite motility, invasion, and replication.

### Photoreactive crosslinking reveals numerous contact points in the IMC6 alveolin domain

Along with other alveolin proteins and associated cytoskeletal proteins, IMC6 is believed to provide structural integrity for the parasite by forming a robust network of filaments just underneath the IMC membrane sacs. The prevailing theory is that this SPN is created primarily by protein-protein interactions via the conserved alveolin domains [1,18]. However, direct experimental evidence for this has been lacking, mainly due to the technical difficulty of identifying interactions between detergent-resistant cytoskeletal proteins. To overcome this challenge, we applied UAA photocrosslinking to the IMC6 alveolin domain to directly test whether this domain is involved in mediating the protein-protein interactions.

To determine which residues to mutate into amber stop codons, we relied on secondary structure and residue burial predictions (Fig 6A and S4 Fig) [31,32]. In total, we chose 38 residues spanning the entire alveolin domain. Each amber mutant was constructed with the IMC6^2-290^ sequence as this truncated protein was fully sufficient for localization and its smaller size facilitates the visualization of upshifted crosslinked products by western blot (Fig 5A). The constructs were generated in a complementation vector with a C-terminal 3xHA epitope tag and randomly integrated into parasites equipped with the aminoacyl-tRNA synthetase/tRNA cassettes (Fig 6B). For each strain expressing an amber mutant, we assessed the incorporation of the UAA (+Azi) and the resulting expression of the mutant protein. We found that none of the amber mutants appeared to affect the localization of the protein; the N162* mutant is shown as a representative example (Fig 6C).

**Figure 6.**
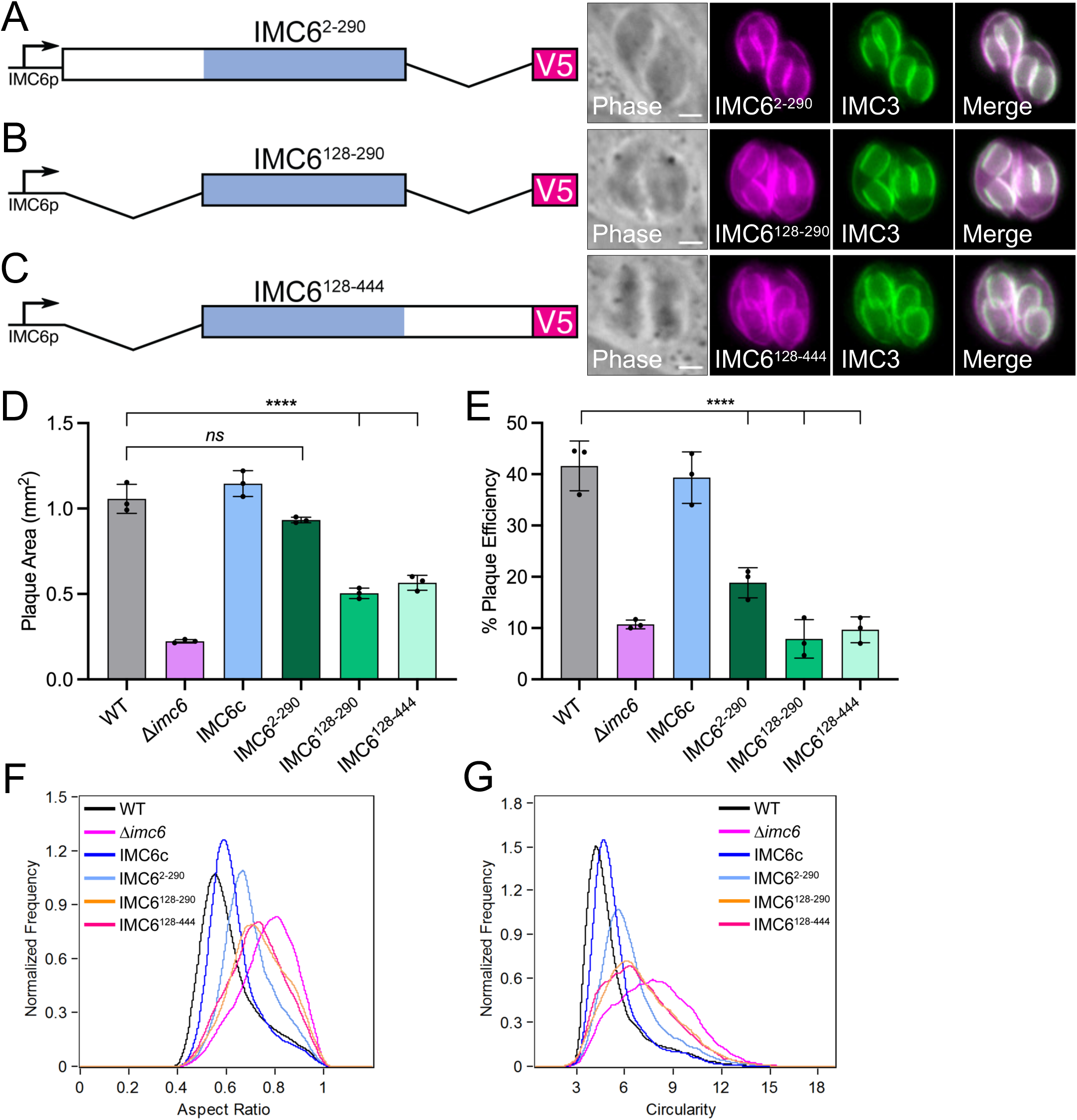
Implementing UAA photoreactive crosslinking for IMC6. (A) Diagram of the IMC6 sequence illustrating the predicted secondary structure. β-sheets are shown in green, the alveolin domain is highlighted in blue, and magenta stars represent residues chosen for crosslinking. (B) Diagram of the complementation construct used for photoreactive crosslinking, which includes the IMC6^2-290^ sequence, driven by the IMC10 promoter with a C-terminal 3xHA epitope tag. (C) IFA of parasites expressing the E2AziRS-3xTy (tRNA synthetase), the Azi-specific tandem tRNA cassettes, and the IMC6^2-290^-3xHA construct with N162 mutated to an amber stop codon. Without Azi added to the system, the N162* mutation results in premature stop in translation and no IMC6-3xHA expression. Upon Azi addition, Azi is incorporated into the amber stop codon and results in proper IMC6-3xHA expression. Magenta, anti-Ty; green, anti-HA. Scale bar is 2 µm.

We divided the amber mutants into five sets of 7 or 8 residues and subjected them to Azi incorporation and photoreactive crosslinking. Upon analyzing each residue by western blot, we observed a myriad of upshifted products throughout the alveolin domain, each representing a crosslink between IMC6 and an unknown binding partner (Fig 7). To discern which upshifts to pursue further, we measured the abundance of crosslinked to uncrosslinked product for every residue and only considered those residues with >10% ratio (S5 Fig). We also excluded residues containing excessive background. Filtering with these criteria resulted in a total of 14 residues. These could be divided into four distinct molecular weights, with each group likely representing a different binding partner. Residues Q185, H203, and F205 were crosslinked at both ∼75 kDa and ∼95 kDa, intriguingly with contrasting intensities (Fig 7A and 7B). Residues N162, Y221, K224, E243, E245, and K277 were crosslinked prominently at ∼175 kDa, and residues E229, Q234, K236, K240, and D249 were crosslinked at ∼140 kDa. Some of these showed upshifts at both ∼140 and ∼175 kDa (K224, E229, Q234, K240, E243, and E245), though with differing intensities. Overall, these crosslinking data demonstrate that the IMC6 alveolin domain contains an abundance of protein-protein interactions with each crosslinked product representing a binding interface.

**Figure 7.**
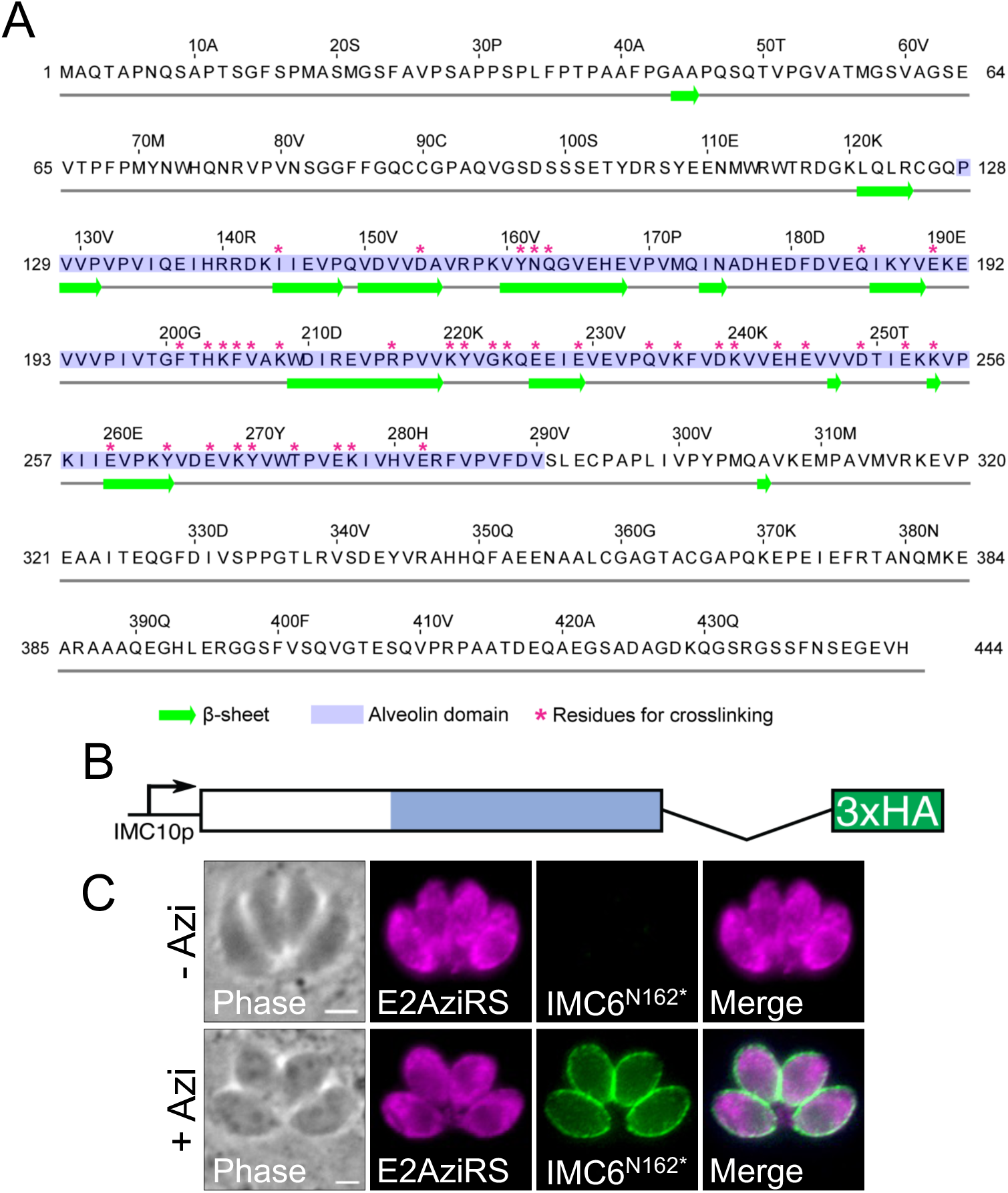
The IMC6 alveolin domain is a major region for protein-protein interactions. (A-E) Western blot of whole cell lysates following Azi incorporation and UV treatment. Black arrowheads represent uncrosslinked IMC6^2-290^-3xHA that migrates at ∼45 kDa. Red stars represent crosslinked upshifts that were chosen for further investigation (ratio of 10% threshold with minimal background, see S1 Table). All blots were detected with anti-HA. (A) Crosslinked upshifts include N162 and Q163 migrating at ∼175 kDa; Q185, E190, and F201 migrating at ∼95 kDa. (B) Crosslinked upshifts include H203 and F205 migrating at ∼75 and ∼95 kDa; V206 and K208 migrating at ∼75 and ∼100 kDa; K220 and Y221 migrating at ∼175 kDa. (C) Crosslinked upshifts include K224, E229, Q234, and K240 migrating at ∼140 and ∼175 kDa; K236 migrating at ∼140 kDa. (D) Crosslinked upshifts include E243 and E245 migrating at ∼140 and ∼175 kDa; D249 migrating at ∼140 kDa; E260 migrating at ∼175 kDa. (E) Crosslinked upshifts include E267, K269, Y270, T273, E276, K277, and E282 migrating at ∼175 kDa, though most of these are extremely faint.

### IMC3 binds IMC6 at multiple residues across the alveolin domain

To identify the crosslinked partners of IMC6, we considered candidates whose localization resembles IMC6 and predicted molecular weight represents the upshifted product. For the larger crosslinked products at ∼140 and ∼175 kDa, IMC3 appeared to be the most likely candidate. To determine if IMC3 is a crosslinked partner of IMC6, we chose N162 to carry out preliminary experiments as proof of concept. We first performed a denaturing co-IP experiment, in which the irradiated and crosslinked samples are boiled in 1% SDS before being diluted to standard RIPA buffer conditions for the IP (Fig 8A) [22]. Probing with anti-HA indicated that we successfully purified both uncrosslinked and crosslinked products, migrating at the same molecular weights as the whole cell lysate samples in Fig 7A. Probing with anti-IMC3 demonstrated that IMC3 is indeed the binding partner for IMC6 at this residue. As a complementary approach, we endogenously tagged IMC3 with a spaghetti monster Myc epitope tag (smMyc) in the amber mutant parasite strain, performed photoreactive crosslinking, and compared the upshift sizes between the untagged and smMyc-tagged samples (Fig 8B). We observed a significant size difference between the untagged and tagged versions, confirming that IMC3 binds IMC6 at residue N162. Based on this preliminary evidence, we evaluated the remaining residues using the endogenous tagging approach as this provides a definitive identity of the binding partner and produces more consistent results than co-IPs.

**Figure 8.**
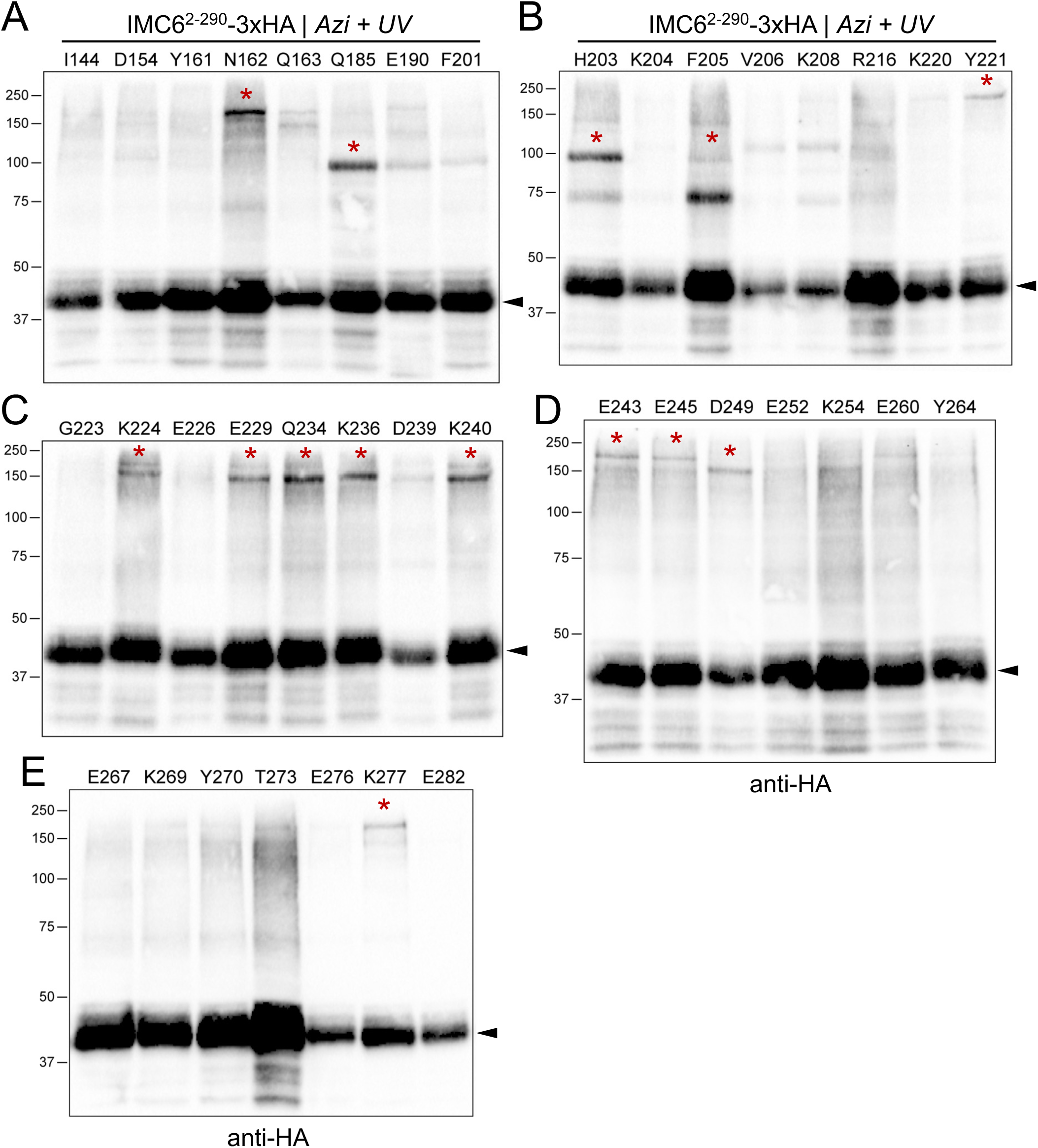
IMC3 binds the IMC6 alveolin domain at multiple sites. (A) Western blot of IMC6-N162* denaturing anti-HA IP following Azi incorporation and UV treatment. Black arrowhead represents uncrosslinked IMC6^2-290^-3xHA (∼45 kDa). Purple arrowhead represents uncrosslinked IMC3 that were nonspecifically captured in the IP (∼85 kDa). Red arrowheads represent the crosslinks-of-interest, seen in both the HA and IMC3 blots (∼175 kDa). The left blot was detected with anti-HA and the right blot was detected with anti-IMC3. (B-L) Western blot of whole cell lysates of the indicated residues following Azi incorporation and UV treatment. For each residue, both wild-type and IMC3-smMyc strains were subjected to photoreactive crosslinking and analyzed on the same blot for direct comparison. Black arrowheads represent uncrosslinked IMC6^2-290^-3xHA (∼45 kDa). Blue arrowheads indicate the crosslinked product at ∼140 kDa, which is not shifted further by the IMC3-smMyc tag. Red arrowheads represent the crosslinks-of-interest, which migrate even higher in the IMC3-smMyc sample (∼175 kDa in the WT vs. ∼200 kDa in the IMC3-smMyc). Orange arrowheads represent *faint* crosslinks-of-interest with similarly higher migration in the IMC3-smMyc sample. Every blot was detected with anti-HA.

We thus endogenously tagged IMC3 with smMyc in the other 10 residues of similar-sized crosslinked products, performed photoreactive crosslinking on both the untagged and tagged versions, and analyzed each pair by western blot (Fig 8C-8L). Five of these residues (Y221, E229, E243, E245, K277) showed an unmistakable size difference due to the smMyc tag, from ∼175 kDa in the untagged version to ∼200 kDa in the tagged version (indicated by red arrowheads). Three other residues (K224, Q234, K240) showed a similar size difference (orange arrowheads). While these almost certainly bind to IMC3 as well, we denoted them as likely IMC3-binding due to the fainter upshifts of the tagged versions. Nonetheless, this demonstrates that IMC3 binds IMC6 via an array of residues spanning the alveolin domain.

We also observed upshifts migrating at ∼140 kDa throughout the alveolin domain (blue arrowheads). These, however, did not shift higher in the IMC3^smMyc^ strain, indicating that IMC3 is not the binding partner for this crosslinked product. While the ∼140 kDa crosslink was the only product for K236 and D249, it was an additional product for all other residues. This suggests that the same residue may have the capacity to interact with both IMC3 and a second, unknown protein or that some background occurs with this shifted product.

### ILP1 is another binding partner of IMC6

We next explored the other three residues with crosslinked upshifts at ∼75 and ∼95 kDa. While the Q185 crosslinked product migrated prominently at ∼95 kDa, the H203 and F205 crosslinked products migrated at both ∼75 and ∼95 kDa, though with opposite intensities (Fig 7A and 7B). We used a similar criteria of protein localization and predicted molecular weight to isolate ILP1 as the prime candidate for these upshifts. Employing the same approach, we endogenously tagged ILP1 with smMyc in each amber mutant parasite strain and performed photoreactive crosslinking on both untagged and tagged strains (Fig 9A-9C). Upon analysis by western blot, we observed a dramatic difference in the migration size between the untagged (∼95 kDa) and smMyc-tagged (∼120 kDa) versions for all three residues. This confirms that ILP1 is indeed the binding partner for IMC6 at these residues. Although the H203 and F205 crosslinked products migrated at two different sizes, only the higher ∼95 kDa was confirmed to bind ILP1 while the target for the lower ∼75 kDa crosslink remains unknown. This indicates that H203 and F205 may bind a second protein in addition to ILP1 or that this product represents background. These confirmed products demonstrate that IMC6 directly binds another cytoskeletal protein via the alveolin domain, underscoring the highly connected nature of the SPN.

**Figure 9.**
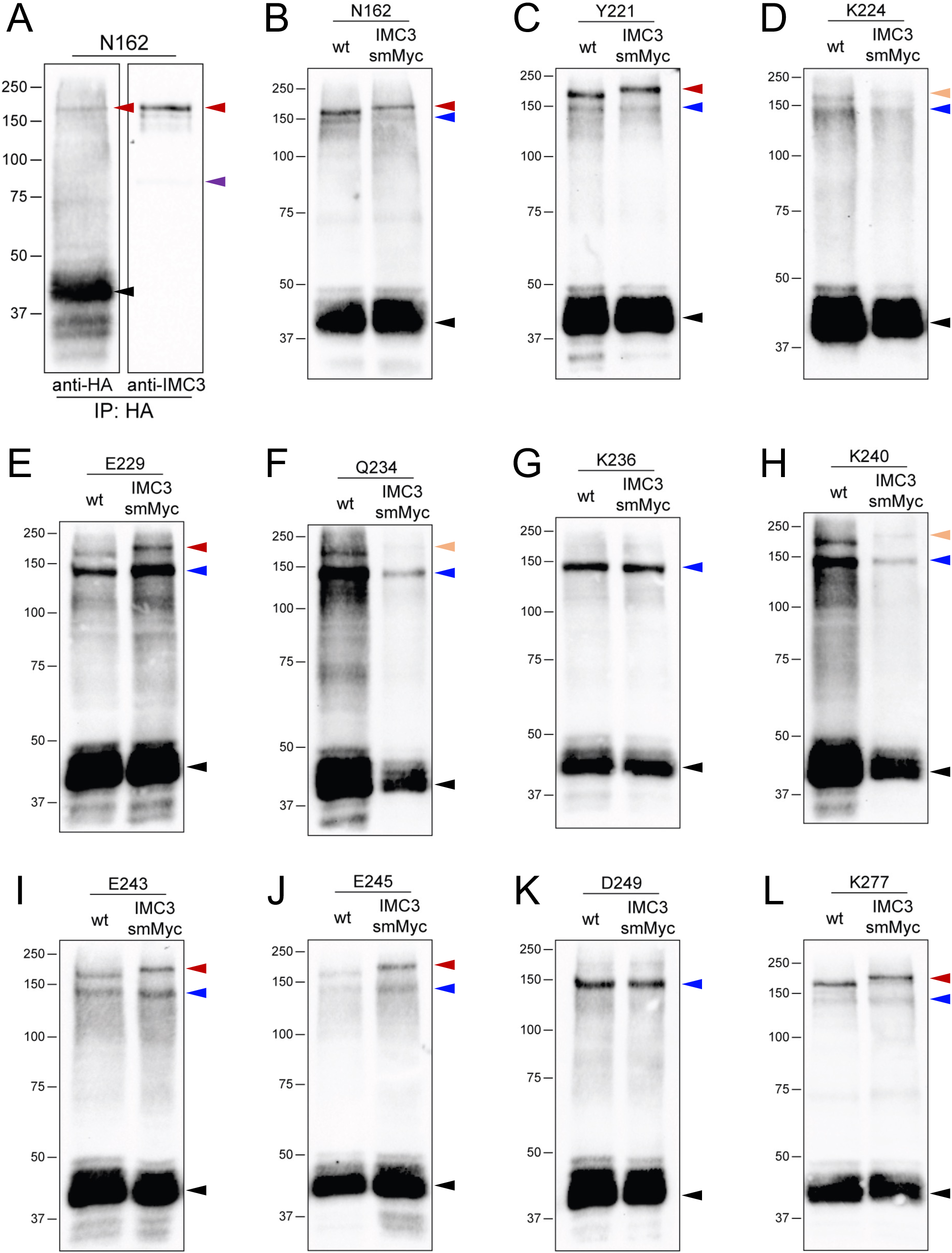
ILP1 binds IMC6 in the alveolin domain. (A-C) Western blot of whole cell lysates of the indicated residues following Azi incorporation and UV treatment. For each residue, both wild-type and ILP1-smMyc strains were subjected to photoreactive crosslinking and analyzed on the same blot for direct comparison. Black arrowheads represent uncrosslinked IMC6^2-290^-3xHA (∼45 kDa). Blue arrowheads indicate the crosslinked product at ∼75 kDa, which is not shifted further by the ILP1-smMyc tag. Red arrowheads represent the crosslinks-of-interest, which migrates even higher in the ILP1-smMyc sample (∼95 kDa vs. ∼120 kDa). Every blot was detected with anti-HA.

## Discussion

By exploring the function and interactions of IMC6, we reveal important insights into the apicomplexan SPN with potentially broader implications for other alveolates. Of the five alveolin proteins that are predicted to be highly fitness-conferring (IMC1, IMC3, IMC4, IMC6, and IMC10), only IMC10 had been examined prior to this study. Conditionally knocking down IMC10 attributed one of the roles of the SPN as a membrane-tether for the mitochondria [33]. Interestingly, however, the knockdown did not affect parasite fitness, though this may be due to the often-incomplete knockdowns of conditional systems. Other alveolins that have been functionally characterized include IMC7, IMC12, and IMC14, which were shown to have modest involvement in tensile strength of extracellular parasites but no growth defects whatsoever [24]. In addition, IMC15 was shown to restrict the number of daughter buds per round of division but again, no growth defects were reported upon its disruption [24]. Thus, this study represents one of the first major perturbations of the *T. gondii* SPN, revealing that it is a critical structure that provides the parasite its elongated shape as well as a scaffold during replication.

Moreover, we clearly link the parasite cytoskeleton to its shape, motility, and invasion. This phenomenon has been reported several times in the past. For example, disruption of TgPhIL1 resulted in shorter and wider parasites with significant motility defects but surprisingly no invasion defects [30,34]. On the other hand, knocking out TgCBAP/TgSIP1 yielded parasites with less severe shape defects but did cause motility and invasion defects [35,36]. We show here that the IMC6 knockout resulted in almost completely rounded parasites, demonstrating that the integrity of the SPN is a key determinant for maintaining parasite shape. We were surprised, though, that this extreme morphological defect did not cause a more severe phenotype in motility and invasion, as both processes were only modestly affected. It is possible that these somewhat subtle defects *in vitro* may be more dramatic *in vivo*, but this requires further study.

Beyond *T. gondii*, some of the *P. berghei* alveolins were shown to be involved in maintaining parasite shape and motility in sporozoites and ookinetes, broadening the significance of the SPN to more distant apicomplexan organisms [37–39,25]. It is unknown if these similarities translate to the human malarial parasite *P. falciparum* or if the SPN provides similar functions during the asexual blood stages. The SPN also harbors cytoskeletal proteins that do not possess alveolin domains, one of which is TgILP1 and the orthologous PbG2. Both were shown to be essential in their respective organisms [19,40,20], again highlighting the conserved functions of these cytoskeletal proteins and the SPN as a whole. One major difference between these parasites is the expanded number of cytoskeletal IMC proteins in *T. gondii*. While it is likely that additional *Plasmodium* IMC proteins will be discovered in the future, the expanded proteome of the *T. gondii* IMC mirrors that of the basal complex [41–43]. Thus, despite the similarity in function, the SPN exhibits important differences in protein composition. Investigation of additional TgIMC proteins may provide further insights into the divergence of these organisms.

Historically, alveolin domains have been thought to mediate the formation of intermediate filament-like structures that constitute the SPN. Evidence for this includes the cytoskeletal properties of all alveolin proteins, the highly conserved repeat motifs that suggest structural functions, and the domain analyses of TgIMC3 and TgIMC8 [44,45]. In other apicomplexans, only sparse evidence exists. The alveolin domain of PbIMC1h was shown to be required for the protein’s localization, and expressing a synthetic protein with alveolin-like motifs in *Tetrahymena* was remarkably sufficient for associating with cytoskeletal structures [25,46]. Here, we demonstrate that the IMC6 alveolin domain binds directly to another alveolin, IMC3, using multiple contact sites. These sites of interaction span almost the entirety of the alveolin domain suggesting the two proteins are highly interwoven. This extensive binding likely increases the strength and avidity of their interaction and ultimately leads to the high tensile strength of filaments. While this method only maps binding interfaces on the bait protein, it is likely that similar experiments using IMC3 would also show this highly interconnected pattern, as the IMC3 alveolin domain is sufficient for trafficking to the SPN [18].

We also demonstrate that ILP1 binds to the alveolin domain of IMC6. Unlike IMC3, however, ILP1 only binds to IMC6 in a 20-residue region of the alveolin domain (aa185-205). In addition, we have previously shown that the variable N-terminal region of IMC6 is required for crosslinking to ILP1 [1]. It is thus possible that ILP1 binds to both regions of IMC6 or that the N-terminus indirectly facilitates ILP1 binding to the alveolin domain. This region also contains two predicted palmitoylated residues (C89 and C90) that may help IMC6 tether to the IMC membranes and optimally incorporate into the SPN. Regardless, this study confirms that ILP1 forms tight interactions with alveolin proteins, highlighting the importance of alveolin-associated proteins like ILP1 as they are also critical for the organization and function of the SPN. As IMC6, IMC3, and ILP1 share an identical localization pattern with basal levels in the maternal IMC with enrichment in the daughter IMC, they appear to form an essential group of “daughter enriched” subfilaments. IMC10 also shares this localization pattern but we have yet to determine how it links to the other three highlighted in our study. Another likely pairing of alveolins are IMC1 and IMC4 which are distributed more equally between the maternal and daughter IMC. Investigating the interactions of this pair using the UAA system may reveal a distinct subfilament. Together, our results corroborate the long-held hypothesis that the alveolin domains are key mediators of SPN architecture and provide a glimpse into the binding interfaces that link these filaments to one another. It will be interesting to dissect which interactions are essential for SPN integrity and to compare our results to future crystallographic or cryoET studies that promise to reveal the entire organization of the network.

## Materials and Methods

### T. gondii culture

Parental *T. gondii* RHΔ*hxgprt*Δ*ku80* (wild-type) and subsequent strains were grown on confluent monolayers of human foreskin fibroblasts (BJ, ATCC, Manassas, VA) at 37°C and 5% CO_2_ in Dulbecco’s Modified Eagle Medium (DMEM) supplemented with 5% fetal bovine serum (Gibco), 5% Cosmic calf serum (Hyclone), and 1x penicillin-streptomycin-L-glutamine (Gibco). For the motility experiments, HFFs and parasites were cultured in 10% v/v heat-inactivated FBS. Drug selections were performed using 1 μM pyrimethamine (dihydrofolate reductase-thymidylate synthase [DHFR-TS]), 50 μg/mL mycophenolic acid-xanthine (HXGPRT), or 40 μM chloramphenicol (CAT) [47–49]. Homologous recombination to the UPRT locus was negatively selected using 5 μM 5-fluorodeoxyuridine (FUDR) [50].

### PCR and plasmid construction

#### Knockout and complementation of *IMC6*

To knock out *IMC6*, the protospacer was designed to target the coding region of IMC6 (TGGT1_220270) and ligated into the pU6-Universal plasmid using P1-2 [51]. The homology-directed repair (HDR) template included 40 bp of homology immediately upstream of the start codon and 40 bp of homology approximately 200bp downstream of the stop codon (P3-4). The HDR template was PCR amplified from a pJET vector containing the HXGPRT selectable marker driven by the NcGRA7 promoter. For transfection, approximately 50 µg of the gRNA-carrying pU6-Universal plasmid was precipitated in ethanol, and the PCR-amplified HDR template was purified by phenol-chloroform extraction and precipitated in ethanol. Both constructs were electroporated into the RHΔ*hxgprt*Δ*ku80* (wild-type) parasite strain. Transfected cells were allowed to invade a confluent monolayer of HFFs overnight, and appropriate selection was subsequently applied. Successful knockout was confirmed by IFA, and clonal lines were obtained through limiting dilution.

All IMC6 complementation constructs were modified from those used in [22]. Each construct contains either the full-length or truncated version of the IMC6 coding sequence, a C-terminal V5 tag, and UPRT homology regions to drive this cassette into the *UPRT* locus. For this study, the ILP1 promoter was replaced with the IMC6 promoter in each plasmid using Gibson assembly. The IMC6 promoter was amplified from genomic DNA (P5-6). The full-length IMC6 coding sequence along with the rest of the plasmid was amplified using P7/P9. These amplicons were purified and ligated using the NEBuilder HiFi DNA Assembly kit to generate the final plasmid. The same process was followed for each truncation: IMC6^2-290^ was amplified using P7/P9, IMC6^128-290^ was amplified using P8/P9, and IMC6^128-444^ was amplified using P8/P9. 100 µg of each complementation plasmid was linearized with DraIII-HF and transfected into Δ*imc6* parasites along with a pU6-gRNA that targets the UPRT coding region. Selection was performed with FUDR for replacement of *UPRT* [50] and potential clones were screened by IFA.

#### Amber stop codon mutagenesis

The plasmid used to express the amber stop codon tRNA and its orthogonal tRNA synthetase is described in [22]. To generate amber mutants, we first created the IMC6 expression construct, which contains the IMC10 promoter driving the coding sequence of IMC6^2-290^ with a C-terminal 3xHA epitope tag as well as the DHFR-TS selectable marker. We then used the NEB Q5 mutagenesis kit to exchange the corresponding residue with the amber stop codon TAG using primers P10-P85. 100 µg of each amber mutant plasmid was linearized with DraIII-HF, transfected into wild-type parasites stably expressing pGra-E2AziTy.HPT.3xtRNA, and selected with pyrimethamine.

#### Endogenous epitope tagging

All genes were C-terminally tagged in this study. A pU6-Universal plasmid was generated containing a protospacer against the 3′ UTR of the target protein approximately 100 bp downstream of the stop codon. The HDR template was PCR amplified using the Δ*ku80*-dependent LIC vector pSmMyc.LIC-CAT, which includes the epitope tag, 3′ UTR, and the selection cassette. The 60-bp primers include 40 bp of homology immediately upstream of the stop codon or 40 bp of homology within the 3′ UTR downstream of the CRISPR/Cas9 cut site. Primers P86-105 were used to generate each of these constructs.

### Antibodies

The HA epitope was detected with mouse monoclonal antibody (mAb) HA.11 (BioLegend; 901515) or rabbit polyclonal antibody (pAb) anti-HA (Invitrogen; PI715500). The Ty1 epitope was detected with mouse mAb BB2 [52]. The V5 epitope was detected with mouse mAb anti-V5 (Invitrogen; R96025). *Toxoplasma*-specific antibodies include mAb mouse anti-IMC1 [53], pAb rabbit anti-IMC3 [16], pAb rabbit anti-IMC6 [22], pAb rat anti-IMC10 [33], pAb rabbit anti-IMC12 [54], mouse mAb anti-ISP1 [55], mAb mouse anti-MIC2 [56], mAb mouse anti-ROP7 [57], pAb rabbit anti-ROP13 [58], pAb rat anti-GRA39 [59], mAb mouse anti-F_1_β subunit (5F4) [60], mAb mouse anti-ATrx1 (11G8) [61], pAb guinea pig anti-NHE3 [62], mAb mouse anti-SAG1 (DG52) [63].

### Immunofluorescence assay and western blot

For IFAs, HFF cells were grown to confluence on glass coverslips. 18–36 hours post infection with *T. gondii*, the coverslips were fixed with 3.7% formaldehyde in PBS and processed for immunofluorescence as described [58]. Primary antibodies were typically incubated for 1 hour, washed with PBS 3-5 times, species-specific secondary antibodies (Alexa Fluor 488/594) were incubated for 1 hour, and washed with PBS at least 3-5 times. Coverslips were then mounted in Vectashield (Vector Labs, Burlingame, CA), viewed with an Axio Imager.Z1 fluorescent microscope (Zeiss), and processed with ZEN 2.3 software (Zeiss).

For western blot, whole cell parasite lysates were prepared in 1x Laemmli sample buffer with 100 mM DTT and boiled at 100°C for 10 minutes. Lysates were resolved by SDS-PAGE and transferred to nitrocellulose membranes. Membranes were incubated in primary antibodies for 1 hour, washed in PBS with 0.01% TWEEN-20, incubated in secondary antibodies conjugated to horseradish peroxidase for 1 hour, and washed with PBS-TWEEN. Chemiluminescence was induced using the SuperSignal West Pico substrate (Pierce) and imaged on a ChemiDoc XRS+ (Bio-Rad, Hercules, CA).

### Parasite functional assays

#### Plaque assay

HFF monolayers were grown in 6-well plates to confluence and subsequently infected with 200 parasites/well and allowed to form plaques for 7 days. Cells were then fixed with ice-cold methanol and stained with crystal violet. The areas of 30 plaques/well were measured using ZEN software (Zeiss) and Fiji. Plaque efficiency was measured by counting the number of plaques/well divided by the number of parasites inoculated/well. Triplicate experiments were conducted for all plaque assays. Graphical and statistical analyses were performed using Prism GraphPad 8.0. Significance of plaque areas was determined by two-way ANOVA and significance of plaque efficiency was determined by multiple two-tailed t-tests.

#### Invasion assay

Invasion assays were modified from previous protocols [64]. Briefly, parasites were mechanically released through a syringe, resuspended in Endo buffer and settled onto coverslips with non-confluent HFF monolayers for 20 min. Endo buffer was then replaced with warm D1 media (Dulbecco’s modified Eagle’s medium, 20 mM HEPES, 1% fetal bovine serum) and incubated at 37 °C for the appropriate invasion time points (5 min, 20 min, 60 min). Coverslips were then fixed and blocked (3% BSA in PBS), and extracellular parasites were stained with anti-SAG1 antibodies. Coverslips were then permeabilized (3% BSA, 0.2% Triton X-100 in PBS), and all parasites were stained with anti-F_1_β ATPase antibody and subsequently the secondary antibodies. Host nuclei were stained with Hoechst during the final rounds of PBS washing. Parasites were scored as invaded (SAG1-, F_1_β+) or not (SAG1+, F_1_β+) per host nucleus. These assays were performed in triplicate with at least 25 host nuclei across 7 fields for each replicate. Significance was determined using two-way ANOVA.

#### Replication defects by IFA

Confluent HFF monolayers on glass coverslips were infected at low MOI, incubated for 1 hour at 37°C, and extracellular parasites were washed away by media changes. Then, 24-and 32-hours post-infection, coverslips were fixed with 3.7% PFA, processed for immunofluorescence, and labeled with anti-ISP1 and anti-IMC3. To score normal vs. abnormal vacuoles, >300 vacuoles across at least 15 fields were counted. Vacuoles were categorized as abnormal if any one of the replication defects shown in Fig 4 were present in a vacuole. All IFAs were performed in triplicate and significance was determined using multiple two-tailed t-tests.

To stain extracellular parasites, wild-type, Δ*imc6*, and IMC6c strains were allowed to egress naturally, collected, and washed in PBS before being settled onto coverslips coated with poly-L-lysine (Sigma Aldrich). Standard IFA procedure was followed to stain samples with mouse anti-ISP2 and rabbit anti-IMC3, making sure to be extra gentle with all wash steps to avoid disturbing the settled parasites. Quantifications were performed by counting >300 parasites across at least 15 fields. Three replicates were conducted, and significance was determined by multiple two-tailed t tests.

### ImageStream flow cytometry

Freshly lysed (or mechanically released) extracellular parasites were processed for immunofluorescence in solution. Parasites were fixed with 3.7% formaldehyde, permeabilized, blocked, and stained with mouse anti-IMC1 followed by anti-mouse Alexa Fluor 488. For each parasite strain, at least 20,000 individual images were acquired using the INSPIRE acquisition software of the ImageStream^®X^ MKII (Cytek Biosciences, Seattle, WA) imaging flow cytometer with 60× magnification and a core width of 6 μM.

All analysis was performed using IDEAS software v6.3.26. The Circularity feature measures the radial variance from the central point of the object. Higher values equate to low radial variance and indicate a highly circular object, while lower values indicate that the object has a high amount of radial variance and have a less circular morphology. The Elongated ML classifier attempts to identify objects that have an elongated shape and assigns a value to each object, indicating confidence. The more positive the value is, the more likely the object is to be elongated. Negative values represent poor prediction confidence, and it is more likely that these objects are circular rather than elongated.

## 3D motility assay

Parasites were mechanically released by syringe lysis and filtered through a 3-μm Whatman Nucleopore filter (Milipore Sigma). Parasites were then centrifuged and resuspended at 5×10^8^ parasites/mL in live cell imaging solution (LCIS) (155 mM NaCl, 3 mM KCl, 2 mM CaCl_2_, 1 mM MgCl_2_, 3 mM NaH_2_PO_4_, 10 mM HEPES, and 20 mM glucose (Sigma-Aldrich) with 20 µg/mL Hoechst 33342 to visualize the nuclei. 1:3:3 volumes of parasites:LCIS:Matrigel were mixed gently, in that order, and pipetted into a Pitta chamber [30]. Flow cells were incubated for 7 min at 27°C and then 3 min at 35°C before imaging. Images were captured using a Nikon Eclipse TE300 epifluorescence microscope (Nikon Instruments, Melville, NY) equipped with a 20X Plan Apo λ objective and a NanoScanZ piezo Z stage insert (Prior Scientific, Rockalnd, MA). 1024 pixel × 384 pixel fluorescence images were captured with an iXON Life 888 EMCCD camera (Andor Technology) driven by NIS Elements software v.5.11 (Nikon Instruments). Fluorescence excitation was controlled using a pE-4000 LED illumination system (CoolLED, Andover England). Images were captured for 80 sec. The camera was set to trigger mode, no binning, readout speed of 30 Mhz, conversion gain of 3.8x, and an EM gain of 300. Image stacks consisted of 41 x-y slices captured 1 μm apart in z for 16 ms. 1024 × 1024 pixel brightfield images were captured over 5 min using a 60X Plan Apo λ objective. Brightfield image stacks consisted of 21 x-y slices captured 1 μm apart in z for 40 ms.

Trajectory data were analyzed with Imaris ×64 v. 9.9.0 software (Bitplane AG, Zurich, Switzerland). Parasites were tracked using an estimated spot volume of 3.0 × 3.0 × 6.0 µm. Tracks were gated for 10s duration and 2.8 µm track displacement in order to be considered moving. The 2.8 µm minimum displacement parameter was determined by both heat-killing and treating all parasite strains with cytochalasin D (CD). *T. gondii* tachyzoites were heat-killed by incubating at 56°C for 30 min. For CD treatment, parasites were incubated in LCIS containing 1 µM CD and 20 µg/mL Hoechst 33342 for a minimum of 20 min and maintaining a concentration of 1 µM CD within the flow cells. All 3D trajectory data were acquired from three biological replicates, each consisting of three technical replicates.

### UAA photoreactive crosslinking

#### Whole cell lysates

All parasites used for photoreactive crosslinking stably express the synthetase/tRNA cassette and IMC6^2-290^ with the appropriate amber mutation. These parasites were allowed to infect confluent HFFs for 16-24 hours, after which the media was replaced with complete DMEM supplemented with 1 mM Azi. After 24 hours of incubation in Azi media, freshly lysed (or syringe-lysed) parasites were collected, resuspended in 2 ml PBS and deposited into 6-well plates. The plates were floated on an iced water slurry without lids and irradiated for 20 minutes in a Spectrolinker XL-1000 UV crosslinker with 365-nm (UV-A) bulbs. Cells were collected by centrifugation and lysed directly in 80-200 µl of sample buffer for SDS-PAGE and subsequent western blot.

For quantification, the band intensities of each primary crosslinked product and each uncrosslinked IMC6^2-290^ were measured using ImageLab software (BioRad). These values were then normalized to background intensities. The ratio was determined by dividing the crosslinked product by the corresponding uncrosslinked product for each applicable lane.

#### Denaturing co-IP

This protocol was followed very closely as previously described [22]. Briefly, following irradiation, parasites were lysed in an extremely harsh SDS buffer (1% SDS, 150 mM NaCl, 50 mM Tris, pH 8.0) and boiled at 100°C for 10 minutes. This lysate was then centrifuged, and the supernatant was diluted 10-fold to RIPA conditions (50 mM Tris, 150 mM NaCl, 0.1% SDS, 0.5% NP-40, 0.5% DOC). IP was then conducted using anti-HA agarose beads (Roche) overnight at 4°C. Affinity captured proteins were eluted directly in sample buffer, agarose beads removed by centrifugation, boiled for 10 minutes, and analyzed on SDS-PAGE.

### Bioinformatic analysis of residue burial predictions

The full IMC6 coding sequence was analyzed by the JPred4 server (http://www.compbio.dundee.ac.uk/jpred4) [31]. This predicts relative solvent accessibility for each residue as well as the secondary structure. In parallel, we also used the Weighted Ensemble Solvent Accessibility (WESA) server, inputting the entire IMC6 coding sequence (https://pipe.rcc.fsu.edu/wesa.html) [32]. This yielded the values in S1 Text. Note that the AlphaFold prediction of TgIMC6 has extremely low confidence values, preventing us from relying on its structure to inform our residue choices. Regardless, the combination of these tools guided our decisions on which residues to mutate in amber stop codons for photoreactive crosslinking.

### Mouse virulence assays

Large vacuoles of RHΔ*hxgprt* (wild-type), Δ*imc6*, IMC6c parasites were syringe-lysed and resuspended in Opti-MEM medium (Thermo Fisher Scientific) prior to intraperitoneal injection into female C57BL/6 mice (4 mice per parasite strain) at the appropriate dosages. An aliquot of this parasite resuspension was used for plaque assays to assess the viability of each injected parasite strain. Mice were monitored for symptoms of infection, weight loss, and survival for 30 days. Survival graphs were generated on Prism GraphPad 8.0.

### Animal experimentation ethics statement

Our protocol was approved by the UCLA Institutional Animal Care and Use Committee (Chancellor’s Animal Research Committee protocol: 2004–005). Mice were euthanized when the animals reached a moribund state and euthanasia was performed following AVMA guidelines.

## Acknowledgements

We thank Salem Haile and Zoran Galic of the UCLA Flow Cytometry Core for help acquiring ImageStream data. We also thank Matthew Rodrigues from Cytek Biosciences for assistance in analyzing and displaying the ImageStream data. We thank Vern Carruthers for the MIC2 antibodies, Gustavo Arrizabalaga for the NHE3 antibody, and Lloyd Kasper and John Boothroyd for the SAG1 antibody. This work was supported by the NIH grants AI064616 to P.J.B. and AI139201 and AI137767 to G.E.W. P.S.B. was also supported by the Ruth L. Kirschstein National Research Service Award GM007185 and UCLA Molecular Biology Institute (MBI) Whitcome Fellowship. A.K.S. was supported by the NIAID predoctoral training grant T32AI055402.

## Supplemental Figure Legends

**S1 Fig. Features used in the “Elongated” ML classifier and their weight.** The ML classifier was used to identify and separate elongated and circular events in all ImageStream flow cytometry samples.

**S2 Fig. Microneme secretion is unaffected in Δ*imc6* parasites.** Western blot of secreted proteins from WT and Δ*imc6* parasites. The calcium ionophore A23187 was used to induce microneme secretion. MIC2 was detected with anti-MIC2 and the constitutively secreted dense granule protein GRA39 was used as a control, detected with anti-GRA39.

**S3 Fig. The apicoplast, mitochondria, PLV/VAC, micronemes, and rhoptries are unaffected in Δ*imc6* parasites.** (A-E) IFAs of mature and budding Δ*imc6* parasites, showing normal morphology of the indicated organelles. (A) The apicoplast was detected with anti-ATrx1 (magenta). (B) Mitochondria were detected with anti-F_1_β (magenta). (C) The PLV/VAC was detected with anti-NHE3 (magenta). (D) Micronemes were detected with anti-MIC2 (magenta). (D) Rhoptries were detected with anti-ROP7 (magenta). All IFAs were costained with anti-IMC3 (green). All scale bars are 2 μm.

**S1 Video. Representative field of WT 3D motility. S2 Video. Close-up view of typical WT 3D motility. S3 Video. Representative field of Δ*imc6* 3D motility.**

**S4 Video. Close-up view of abnormal Δ*imc6* trajectory 1. S5 Video. Close-up view of abnormal Δ*imc6* trajectory 2. S6 Video. Close-up view of abnormal Δ*imc6* trajectory 3. S7 Video. Representative field of IMC6c 3D motility.**

**S8 Video. Close-up view of restored IMC6c 3D motility.**

**S1 Text. Residue burial prediction by WESA.** Residues chosen for amber mutations are bolded.

**S1 Table. Ratio of crosslinked to uncrosslinked products.** Residues chosen for partner identification are bolded.

**S2 Table. Oligonucleotides used in this study.** All primers are listed in 5’ to 3’ orientation.

