## Supplemental Figures S1-3 for "The *Toxoplasma* subpellicular network is highly interconnected and defines parasite shape for efficient motility and replication"

| Feature | Weight (%) |
| --- | --- |
| H Entropy Mean (Fluorescence) | 12.66 |
| H Entropy Std (Fluorescence) | 12.66 |
| H Entropy Std (Brightfield) | 11.29 |
| Major Axis_Morphology (Fluorescence) | 9.65 |
| H Entropy Mean (Brightfield) | 9.18 |
| H Entropy Std (Brightfield) | 9.02 |
| Length_Morphology (Fluorescence) | 8.96 |
| Height_Morphology (Fluorescence) | 8.96 |
| Aspect Ratio | -8.9 |
| Aspect Ratio Intensity (Fluorescence) | -8.71 |

**S1 Fig. Features used in the "Elongated" ML classifier and their weight.** The ML classifier was used to identify and separate elongated and circular events in all ImageStream flow cytometry samples.

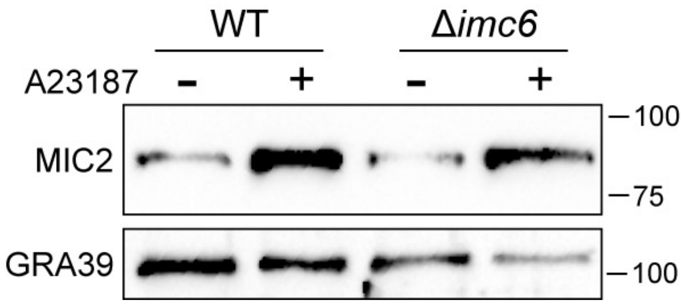


**S2 Fig. Microneme secretion is unaffected in Δ*imc6* parasites.** Western blot of secreted proteins from WT and Δ*imc6* parasites. The calcium ionophore A23187 was used to induce microneme secretion. MIC2 was detected with anti-MIC2 and the constitutively secreted dense granule protein GRA39 was used as a control, detected with anti-GRA39.

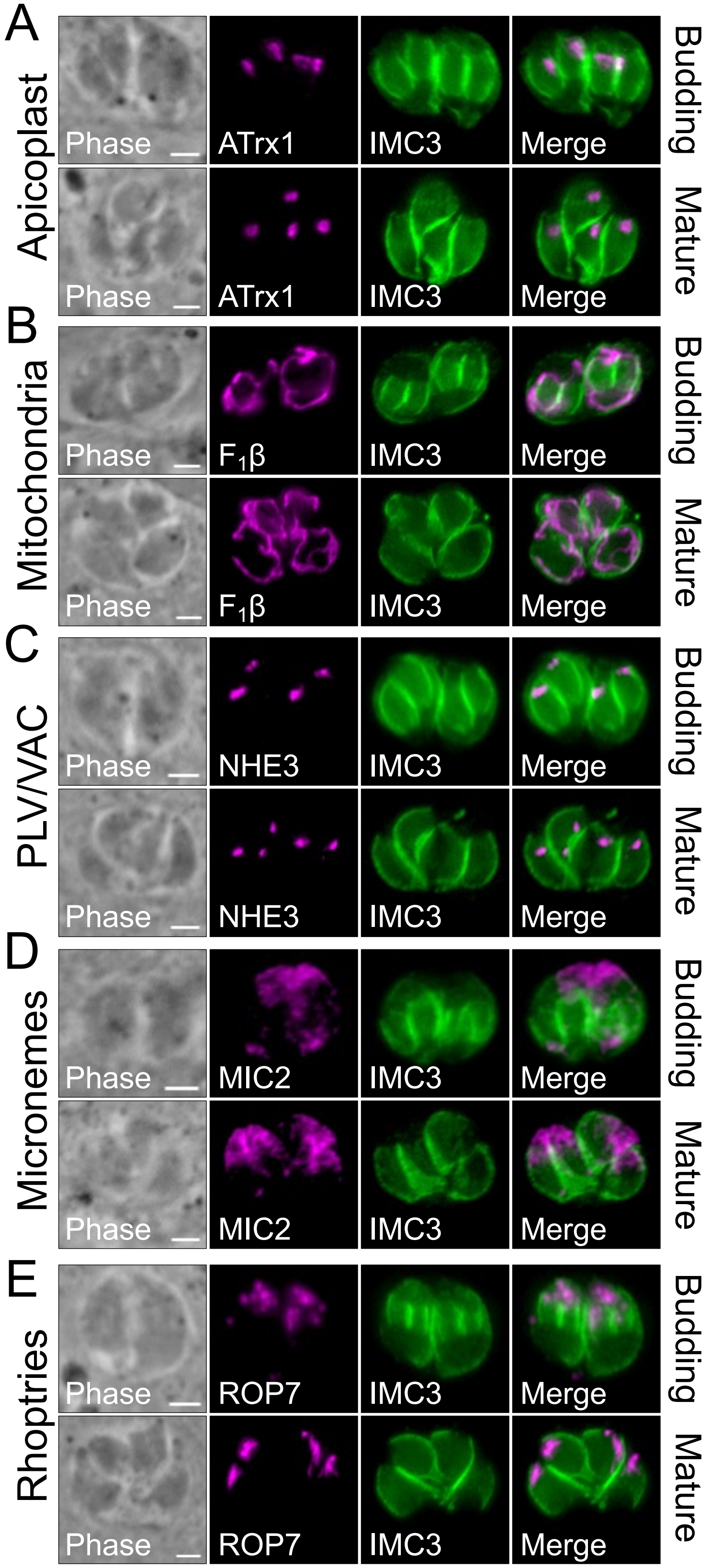
A
Apicoplast
Phase
ATrx1
IMC3
Merge
Budding
Mature
B
Mitochondria
F1β
C
PLV/VAC
NHE3
D
Micronemes
MIC2
E
Rhoptries
ROP7

**S3 Fig. The apicoplast, mitochondria, PLV/VAC, micronemes, and rhoptries are unaffected in Δ*imc6* parasites.** (A-E) IFAs of mature and budding Δ*imc6* parasites, showing normal morphology of the indicated organelles. (A) The apicoplast was detected with anti-ATrx1 (magenta). (B) Mitochondria were detected with anti-$F_1β$ (magenta). (C) The PLV/VAC was detected with anti-NHE3 (magenta). (D) Micronemes were detected with anti-MIC2 (magenta). (D) Rhoptries were detected with anti-ROP7 (magenta). All IFAs were costained with anti-IMC3 (green). All scale bars are 2 μm.
