## Supplemental Text S1 for "The *Toxoplasma* subpellicular network is highly interconnected and defines parasite shape for efficient motility and replication"

**Prediction by WESA - Weighted Ensemble Solvent Accessibility predictor**

Column 1: Residue number
Column 2: One-letter amino acid code

BS : Bayesian Statistics
MLR : Multiple Linear Regression
DT : Decision Tree
NN : Neural Network
SVM : Support Vector Machine
WE : Weighted Ensemble

Col. 3-8: 1 = Exposed; 0 = Buried; [...] = Confidence

***Residues chosen for amber mutations in this study are bolded.***

| | | BS | MLR | DT | NN | SVM | WE |
|---|---|---|---|---|---|---|---|
| 1 | M | 0[0.62] | 1[0.45] | 1[0.84] | 1[0.94] | 1[0.84] | 1[0.73] |
| 2 | A | 0[0.62] | 1[0.45] | 1[0.80] | 1[0.70] | 1[0.88] | 1[0.66] |
| 3 | Q | 1[0.84] | 1[0.87] | 1[0.84] | 1[0.76] | 1[0.94] | 1[0.79] |
| 4 | T | 1[0.38] | 1[0.71] | 1[0.64] | 1[0.58] | 1[0.77] | 1[0.60] |
| 5 | A | 0[0.56] | 1[0.33] | 1[0.58] | 1[0.22] | 1[0.52] | 1[0.30] |
| 6 | P | 0[0.10] | 1[0.87] | 1[0.64] | 1[0.44] | 1[0.76] | 1[0.54] |
| 7 | N | 1[0.50] | 1[0.86] | 1[1.00] | 1[0.76] | 1[0.88] | 1[0.78] |
| 8 | Q | 1[0.48] | 1[0.80] | 1[0.84] | 1[0.76] | 1[0.84] | 1[0.73] |
| 9 | S | 1[0.10] | 1[0.70] | 1[0.64] | 1[0.68] | 1[0.80] | 1[0.64] |
| 10 | A | 0[0.35] | 1[0.13] | 0[0.44] | 1[0.34] | 0[0.03] | 0[0.01] |
| 11 | P | 0[0.17] | 1[0.59] | 0[0.14] | 1[0.36] | 1[0.12] | 1[0.13] |
| 12 | T | 0[0.03] | 1[0.46] | 1[0.26] | 1[0.44] | 1[0.18] | 1[0.24] |
| 13 | S | 0[0.20] | 1[0.67] | 1[0.82] | 1[0.70] | 1[0.77] | 1[0.65] |
| 14 | G | 0[0.53] | 1[0.22] | 0[0.32] | 1[0.00] | 0[0.36] | 0[0.24] |
| 15 | F | 0[0.81] | 0[0.35] | 0[0.40] | 0[0.34] | 0[0.55] | 0[0.51] |
| 16 | S | 0[0.45] | 1[0.25] | 0[0.20] | 1[0.54] | 1[0.47] | 1[0.29] |
| 17 | P | 0[0.60] | 1[0.35] | 0[0.22] | 1[0.06] | 0[0.19] | 0[0.14] |
| 18 | M | 0[0.81] | 0[0.18] | 0[0.60] | 0[0.30] | 0[0.50] | 0[0.49] |
| 19 | A | 0[0.86] | 0[0.26] | 0[0.46] | 0[0.22] | 0[0.29] | 0[0.37] |
| 20 | S | 0[0.80] | 1[0.30] | 0[0.00] | 1[0.38] | 1[0.45] | 1[0.24] |
| 21 | M | 0[0.94] | 0[0.28] | 0[0.82] | 0[0.06] | 0[0.13] | 0[0.31] |
| 22 | G | 0[0.85] | 0[0.12] | 0[0.22] | 0[0.06] | 0[0.37] | 0[0.29] |
| 23 | S | 0[0.83] | 1[0.38] | 1[0.02] | 1[0.18] | 1[0.38] | 1[0.15] |
| 24 | F | 0[0.95] | 0[0.47] | 0[0.64] | 0[0.24] | 0[0.65] | 0[0.55] |
| 25 | A | 0[0.92] | 0[0.26] | 0[0.84] | 0[0.24] | 0[0.44] | 0[0.49] |
| 26 | V | 0[0.89] | 0[0.43] | 0[0.46] | 0[0.50] | 0[0.60] | 0[0.60] |
| 27 | P | 0[0.78] | 1[0.61] | 1[0.24] | 1[0.24] | 0[0.15] | 1[0.03] |
| 28 | S | 0[0.69] | 1[0.66] | 1[0.40] | 1[0.64] | 1[0.60] | 1[0.48] |
| 29 | A | 0[0.89] | 1[0.29] | 0[0.48] | 1[0.06] | 0[0.24] | 0[0.21] |
| 30 | P | 0[0.85] | 1[0.71] | 1[0.02] | 1[0.40] | 0[0.08] | 1[0.08] |
| 31 | P | 0[0.73] | 1[0.72] | 0[0.16] | 1[0.12] | 1[0.08] | 1[0.02] |
| 32 | S | 0[0.56] | 1[0.57] | 1[0.00] | 1[0.60] | 1[0.43] | 1[0.34] |
| 33 | P | 0[0.66] | 1[0.65] | 0[0.16] | 1[0.32] | 0[0.02] | 1[0.05] |
| 34 | L | 0[0.87] | 0[0.09] | 0[0.48] | 0[0.10] | 0[0.54] | 0[0.41] |
| 35 | F | 0[0.94] | 0[0.48] | 0[0.26] | 0[0.32] | 0[0.58] | 0[0.50] |
| 36 | P | 0[0.84] | 1[0.50] | 0[0.20] | 1[0.32] | 0[0.18] | 0[0.03] |
| 37 | T | 0[0.82] | 1[0.01] | 1[0.42] | 0[0.08] | 0[0.32] | 0[0.18] |
| 38 | P | 0[0.92] | 1[0.33] | 1[0.24] | 1[0.10] | 0[0.23] | 0[0.08] |

| | | | | | | | |
|---|---|---|---|---|---|---|---|
| 39 | A | 0[0.98] | 1[0.09] | 0[0.10] | 0[0.20] | 0[0.33] | 0[0.30] |
| 40 | A | 0[0.98] | 1[0.07] | 0[0.28] | 1[0.10] | 0[0.21] | 0[0.17] |
| 41 | F | 0[0.92] | 0[0.63] | 0[0.82] | 0[0.38] | 0[0.73] | 0[0.67] |
| 42 | P | 0[0.79] | 1[0.61] | 1[0.20] | 1[0.08] | 0[0.18] | 0[0.05] |
| 43 | G | 0[0.72] | 1[0.50] | 1[0.42] | 1[0.06] | 0[0.04] | 1[0.02] |
| 44 | A | 0[0.82] | 1[0.15] | 0[0.26] | 1[0.16] | 1[0.01] | 0[0.06] |
| 45 | A | 0[0.71] | 1[0.07] | 0[0.04] | 1[0.12] | 1[0.03] | 0[0.03] |
| 46 | P | 0[0.20] | 1[0.72] | 1[0.22] | 1[0.26] | 1[0.47] | 1[0.29] |
| 47 | Q | 1[0.37] | 1[0.79] | 1[0.86] | 1[0.70] | 1[0.78] | 1[0.69] |
| 48 | S | 1[0.05] | 1[0.61] | 1[0.02] | 1[0.70] | 1[0.51] | 1[0.44] |
| 49 | Q | 1[0.43] | 1[0.74] | 1[0.50] | 1[0.66] | 1[0.71] | 1[0.59] |
| 50 | T | 0[0.04] | 1[0.35] | 1[0.82] | 1[0.26] | 1[0.28] | 1[0.29] |
| 51 | V | 0[0.71] | 0[0.57] | 0[0.44] | 0[0.48] | 0[0.72] | 0[0.64] |
| 52 | P | 0[0.45] | 1[0.61] | 1[0.22] | 1[0.24] | 1[0.18] | 1[0.16] |
| 53 | G | 0[0.58] | 0[0.10] | 0[0.16] | 0[0.38] | 0[0.62] | 0[0.48] |
| 54 | V | 0[0.85] | 0[0.61] | 0[0.46] | 0[0.38] | 0[0.56] | 0[0.55] |
| 55 | A | 0[0.86] | 0[0.36] | 0[0.24] | 0[0.32] | 0[0.56] | 0[0.48] |
| 56 | T | 0[0.82] | 1[0.10] | 0[0.38] | 1[0.18] | 0[0.15] | 0[0.13] |
| 57 | M | 0[0.93] | 0[0.52] | 0[0.64] | 0[0.16] | 0[0.62] | 0[0.52] |
| 58 | G | 0[0.76] | 0[0.21] | 0[0.22] | 0[0.22] | 0[0.42] | 0[0.37] |
| 59 | S | 0[0.56] | 1[0.21] | 0[0.20] | 1[0.30] | 1[0.38] | 1[0.16] |
| 60 | V | 0[0.76] | 0[0.37] | 0[0.26] | 1[0.00] | 0[0.60] | 0[0.38] |
| 61 | A | 0[0.65] | 0[0.07] | 0[0.10] | 1[0.16] | 1[0.13] | 1[0.00] |
| 62 | G | 0[0.45] | 1[0.28] | 1[0.40] | 1[0.44] | 1[0.28] | 1[0.27] |
| 63 | S | 0[0.20] | 1[0.55] | 0[0.38] | 1[0.72] | 1[0.67] | 1[0.43] |
| 64 | E | 1[0.01] | 1[0.81] | 1[1.00] | 1[0.74] | 1[0.70] | 1[0.68] |
| 65 | V | 0[0.71] | 0[0.36] | 0[0.08] | 0[0.40] | 0[0.46] | 0[0.45] |
| 66 | T | 0[0.26] | 1[0.01] | 0[0.14] | 1[0.12] | 0[0.13] | 0[0.09] |
| 67 | P | 0[0.35] | 1[0.49] | 1[0.04] | 1[0.22] | 0[0.01] | 1[0.05] |
| 68 | F | 0[0.56] | 0[0.61] | 0[0.82] | 0[0.20] | 0[0.61] | 0[0.55] |
| 69 | P | 0[0.41] | 1[0.10] | 1[0.42] | 0[0.16] | 0[0.25] | 0[0.16] |
| 70 | M | 0[0.52] | 0[0.32] | 0[0.42] | 0[0.32] | 0[0.48] | 0[0.46] |
| 71 | Y | 0[0.41] | 0[0.35] | 0[0.44] | 0[0.08] | 0[0.42] | 0[0.35] |
| 72 | N | 1[0.29] | 1[0.47] | 1[0.00] | 1[0.14] | 1[0.13] | 1[0.08] |
| 73 | W | 0[0.50] | 0[0.40] | 0[0.52] | 0[0.40] | 0[0.59] | 0[0.55] |
| 74 | H | 1[0.20] | 1[0.45] | 0[0.24] | 1[0.20] | 1[0.26] | 1[0.11] |
| 75 | Q | 1[0.40] | 1[0.69] | 1[0.34] | 1[0.72] | 1[0.81] | 1[0.62] |
| 76 | N | 1[0.39] | 1[0.69] | 1[0.22] | 1[0.68] | 1[0.72] | 1[0.56] |
| 77 | R | 1[0.36] | 1[0.77] | 1[0.48] | 1[0.72] | 1[0.66] | 1[0.59] |
| 78 | V | 0[0.37] | 0[0.35] | 0[0.26] | 0[0.50] | 0[0.47] | 0[0.50] |
| 79 | P | 0[0.05] | 1[0.52] | 1[0.28] | 1[0.22] | 0[0.03] | 1[0.09] |
| 80 | V | 0[0.56] | 0[0.37] | 0[0.38] | 0[0.48] | 0[0.26] | 0[0.44] |
| 81 | N | 1[0.14] | 1[0.63] | 1[0.24] | 1[0.48] | 1[0.51] | 1[0.39] |
| 82 | S | 0[0.27] | 1[0.24] | 0[0.02] | 1[0.34] | 0[0.02] | 1[0.06] |
| 83 | G | 0[0.43] | 1[0.06] | 0[0.20] | 0[0.16] | 0[0.35] | 0[0.29] |
| 84 | G | 0[0.45] | 1[0.06] | 0[0.60] | 0[0.28] | 0[0.26] | 0[0.36] |
| 85 | F | 0[0.66] | 0[0.52] | 0[0.42] | 0[0.64] | 0[0.59] | 0[0.64] |
| 86 | F | 0[0.71] | 0[0.69] | 0[0.84] | 0[0.60] | 0[0.62] | 0[0.71] |
| 87 | G | 0[0.59] | 0[0.38] | 0[0.42] | 0[0.72] | 0[0.60] | 0[0.66] |
| 88 | Q | 0[0.06] | 1[0.50] | 1[0.50] | 0[0.04] | 1[0.19] | 1[0.10] |
| 89 | C | 0[0.71] | 0[0.64] | 0[0.66] | 0[0.78] | 0[0.47] | 0[0.69] |
| 90 | C | 0[0.64] | 0[0.53] | 0[0.82] | 0[0.56] | 0[0.61] | 0[0.68] |
| 91 | G | 0[0.39] | 1[0.01] | 0[0.16] | 0[0.02] | 0[0.37] | 0[0.24] |
| 92 | P | 0[0.28] | 1[0.46] | 1[0.44] | 0[0.16] | 0[0.20] | 0[0.11] |
| 93 | A | 0[0.47] | 0[0.19] | 0[0.32] | 1[0.06] | 0[0.29] | 0[0.22] |
| 94 | Q | 1[0.30] | 1[0.63] | 1[0.84] | 1[0.74] | 1[0.65] | 1[0.64] |
| 95 | V | 0[0.60] | 0[0.38] | 0[0.64] | 0[0.36] | 0[0.53] | 0[0.53] |

| | | | | | | | |
|---|---|---|---|---|---|---|---|
| 96 | G | 1[0.16] | 1[0.28] | 1[0.18] | 1[0.26] | 1[0.22] | 1[0.17] |
| 97 | S | 1[0.29] | 1[0.64] | 0[0.18] | 1[0.62] | 1[0.64] | 1[0.44] |
| 98 | D | 1[0.59] | 1[0.93] | 1[0.46] | 1[0.26] | 1[0.60] | 1[0.42] |
| 99 | S | 1[0.56] | 1[0.69] | 1[0.34] | 1[0.64] | 1[0.59] | 1[0.52] |
| 100 | S | 1[0.64] | 1[0.64] | 1[0.22] | 1[0.36] | 1[0.54] | 1[0.38] |
| 101 | P | 1[0.24] | 1[0.71] | 1[0.42] | 1[0.26] | 1[0.19] | 1[0.23] |
| 102 | E | 1[0.75] | 1[0.90] | 1[1.00] | 1[0.42] | 1[0.73] | 1[0.61] |
| 103 | T | 0[0.16] | 0[0.10] | 1[0.20] | 1[0.36] | 0[0.06] | 1[0.07] |
| 104 | Y | 1[0.41] | 1[0.21] | 1[0.20] | 1[0.20] | 1[0.13] | 1[0.12] |
| 105 | D | 1[0.73] | 1[0.70] | 1[0.44] | 1[0.60] | 1[0.45] | 1[0.47] |
| 106 | R | 1[0.65] | 1[0.65] | 1[1.00] | 1[0.66] | 1[0.56] | 1[0.61] |
| 107 | S | 1[0.56] | 1[0.57] | 1[0.78] | 1[0.84] | 1[0.85] | 1[0.74] |
| 108 | Y | 0[0.46] | 0[0.24] | 0[0.40] | 0[0.50] | 0[0.32] | 0[0.46] |
| 109 | E | 1[0.42] | 1[0.72] | 1[0.84] | 1[0.60] | 1[0.66] | 1[0.60] |
| 110 | E | 1[0.66] | 1[0.58] | 1[0.84] | 1[0.62] | 1[0.69] | 1[0.62] |
| 111 | N | 1[0.58] | 1[0.61] | 1[0.66] | 1[0.76] | 1[0.73] | 1[0.66] |
| 112 | M | 0[0.74] | 0[0.70] | 0[0.66] | 0[0.80] | 0[0.49] | 0[0.71] |
| 113 | W | 0[0.58] | 0[0.56] | 0[0.84] | 0[0.54] | 0[0.65] | 0[0.69] |
| 114 | R | 1[0.60] | 1[0.45] | 1[0.44] | 1[0.70] | 1[0.54] | 1[0.52] |
| 115 | W | 1[0.04] | 0[0.24] | 0[0.56] | 0[0.56] | 0[0.59] | 0[0.59] |
| 116 | T | 0[0.53] | 0[0.53] | 0[0.82] | 0[0.56] | 0[0.71] | 0[0.71] |
| 117 | R | 1[0.46] | 1[0.65] | 1[0.28] | 1[0.48] | 0[0.01] | 1[0.22] |
| 118 | D | 1[0.49] | 1[0.82] | 1[0.46] | 1[0.60] | 1[0.63] | 1[0.54] |
| 119 | G | 1[0.10] | 1[0.40] | 0[0.20] | 0[0.18] | 0[0.31] | 0[0.24] |
| 120 | K | 1[0.21] | 1[0.48] | 1[0.70] | 1[0.30] | 1[0.36] | 1[0.33] |
| 121 | L | 0[0.41] | 0[0.47] | 0[0.22] | 0[0.34] | 0[0.10] | 0[0.30] |
| 122 | Q | 1[0.15] | 1[0.82] | 1[0.84] | 1[0.56] | 1[0.57] | 1[0.55] |
| 123 | L | 0[0.21] | 0[0.35] | 0[1.00] | 0[0.48] | 0[0.64] | 0[0.66] |
| 124 | R | 1[0.30] | 1[0.89] | 1[0.50] | 1[0.50] | 1[0.54] | 1[0.47] |
| 125 | C | 0[0.43] | 0[0.41] | 0[0.64] | 0[0.52] | 0[0.70] | 0[0.65] |
| 126 | G | 1[0.23] | 1[0.42] | 1[0.42] | 1[0.82] | 1[0.56] | 1[0.55] |
| 127 | Q | 1[0.38] | 1[0.79] | 1[0.46] | 1[0.68] | 1[0.70] | 1[0.59] |
| 128 | P | 1[0.11] | 1[0.83] | 1[0.66] | 1[0.74] | 1[0.43] | 1[0.54] |
| 129 | V | 0[0.41] | 0[0.47] | 0[1.00] | 0[0.38] | 0[0.46] | 0[0.57] |
| 130 | V | 0[0.31] | 0[0.52] | 0[1.00] | 0[0.76] | 0[0.72] | 0[0.81] |
| 131 | P | 1[0.22] | 1[0.79] | 1[0.46] | 1[0.78] | 1[0.70] | 1[0.62] |
| 132 | V | 0[0.45] | 0[0.15] | 0[0.32] | 0[0.50] | 0[0.64] | 0[0.56] |
| 133 | P | 0[0.13] | 1[0.25] | 0[0.46] | 1[0.18] | 1[0.16] | 1[0.01] |
| 134 | V | 1[0.36] | 1[0.46] | 0[0.02] | 1[0.78] | 1[0.49] | 1[0.45] |
| 135 | I | 0[0.46] | 0[0.18] | 0[0.30] | 0[0.24] | 0[0.32] | 0[0.34] |
| 136 | Q | 1[0.41] | 1[0.68] | 1[0.64] | 1[0.86] | 1[0.74] | 1[0.69] |
| 137 | E | 1[0.60] | 1[0.72] | 1[1.00] | 1[0.90] | 1[0.78] | 1[0.79] |
| 138 | I | 1[0.35] | 0[0.10] | 1[0.36] | 1[0.02] | 1[0.02] | 1[0.01] |
| 139 | H | 1[0.02] | 1[0.76] | 1[0.64] | 1[0.60] | 1[0.46] | 1[0.49] |
| 140 | R | 1[0.28] | 1[0.58] | 1[0.18] | 1[0.40] | 1[0.22] | 1[0.25] |
| 141 | R | 1[0.46] | 1[0.74] | 1[0.62] | 1[0.64] | 0[0.03] | 1[0.33] |
| 142 | D | 1[0.51] | 1[0.87] | 1[0.46] | 1[0.74] | 1[0.36] | 1[0.50] |
| 143 | K | 0[0.14] | 1[0.65] | 1[0.64] | 0[0.02] | 0[0.05] | 1[0.05] |
| **144** | **I** | **0[0.17]** | **0[0.12]** | **0[1.00]** | **1[0.10]** | **0[0.04]** | **0[0.20]** |
| 145 | I | 0[0.35] | 0[0.15] | 0[1.00] | 0[0.62] | 0[0.56] | 0[0.67] |
| 146 | E | 1[0.24] | 1[0.93] | 1[0.30] | 1[0.88] | 1[0.61] | 1[0.61] |
| 147 | V | 0[0.41] | 0[0.15] | 0[1.00] | 0[0.12] | 0[0.51] | 0[0.47] |
| 148 | P | 0[0.16] | 1[0.49] | 0[0.60] | 0[0.08] | 0[0.47] | 0[0.32] |
| 149 | Q | 1[0.34] | 1[0.46] | 1[0.06] | 1[0.68] | 0[0.10] | 1[0.21] |
| 150 | V | 0[0.22] | 1[0.28] | 0[0.82] | 1[0.20] | 0[0.31] | 0[0.21] |
| 151 | D | 1[0.45] | 1[0.66] | 1[0.66] | 1[0.86] | 1[0.63] | 1[0.65] |
| 152 | V | 0[0.51] | 0[0.30] | 0[0.82] | 1[0.16] | 0[0.35] | 0[0.29] |

| | | | | | | | |
|---|---|---|---|---|---|---|---|
| 153 | V | 1[0.73] | 1[0.43] | 1[0.80] | 1[0.78] | 1[0.52] | 1[0.60] |
| **154** | **D** | **1[0.68]** | **1[0.82]** | **1[0.44]** | **1[0.78]** | **1[0.40]** | **1[0.53]** |
| 155 | A | 1[0.70] | 1[0.61] | 1[0.54] | 1[0.76] | 1[0.50] | 1[0.56] |
| 156 | V | 1[0.35] | 1[0.06] | 0[0.66] | 1[0.18] | 0[0.11] | 0[0.12] |
| 157 | R | 1[0.74] | 1[0.40] | 1[0.64] | 1[0.52] | 1[0.15] | 1[0.34] |
| 158 | P | 1[0.76] | 1[0.81] | 1[0.64] | 1[0.54] | 1[0.54] | 1[0.52] |
| 159 | K | 1[0.91] | 1[0.92] | 1[1.00] | 1[0.84] | 1[0.66] | 1[0.74] |
| 160 | V | 0[0.64] | 1[0.03] | 0[0.08] | 0[0.10] | 0[0.56] | 0[0.34] |
| **161** | **Y** | **1[0.77]** | **1[0.83]** | **0[0.40]** | **1[0.86]** | **1[0.28]** | **1[0.39]** |
| **162** | **N** | **1[0.77]** | **1[0.63]** | **1[0.02]** | **1[0.60]** | **1[0.22]** | **1[0.32]** |
| **163** | **Q** | **1[0.95]** | **1[0.96]** | **1[0.84]** | **1[0.90]** | **1[0.77]** | **1[0.79]** |
| 164 | G | 1[0.96] | 1[0.81] | 1[0.46] | 1[0.92] | 1[0.92] | 1[0.78] |
| 165 | V | 1[0.98] | 1[0.36] | 0[0.40] | 1[0.06] | 0[0.33] | 0[0.16] |
| 166 | E | 1[0.97] | 1[0.93] | 1[1.00] | 1[0.90] | 1[0.87] | 1[0.85] |
| 167 | H | 1[1.00] | 1[0.87] | 1[0.18] | 1[0.82] | 1[0.42] | 1[0.53] |
| 168 | E | 1[1.00] | 1[0.86] | 1[1.00] | 1[0.92] | 1[0.90] | 1[0.86] |
| 169 | V | 1[1.00] | 1[0.28] | 0[0.30] | 0[0.40] | 0[0.52] | 0[0.39] |
| 170 | P | 1[0.99] | 1[0.87] | 1[0.78] | 1[0.90] | 1[0.84] | 1[0.80] |
| 171 | V | 1[0.99] | 1[0.71] | 0[0.44] | 1[0.76] | 1[0.56] | 1[0.45] |
| 172 | M | 1[0.43] | 1[0.01] | 0[0.84] | 0[0.32] | 0[0.50] | 0[0.47] |
| 173 | Q | 1[0.96] | 1[0.57] | 1[0.84] | 1[0.68] | 1[0.59] | 1[0.61] |
| 174 | I | 1[0.97] | 1[0.59] | 0[0.20] | 1[0.80] | 1[0.54] | 1[0.49] |
| 175 | N | 1[0.83] | 1[0.70] | 1[0.12] | 1[0.86] | 1[0.27] | 1[0.46] |
| 176 | A | 0[0.66] | 1[0.09] | 0[0.08] | 1[0.32] | 0[0.45] | 0[0.14] |
| 177 | D | 1[0.71] | 1[0.84] | 1[1.00] | 1[0.88] | 1[0.81] | 1[0.80] |
| 178 | H | 1[0.63] | 1[0.90] | 1[0.84] | 1[0.90] | 1[0.96] | 1[0.84] |
| 179 | E | 1[0.63] | 1[0.91] | 1[1.00] | 1[0.86] | 1[0.71] | 1[0.76] |
| 180 | D | 1[0.64] | 1[0.74] | 1[0.48] | 1[0.92] | 1[0.56] | 1[0.64] |
| 181 | F | 0[0.49] | 1[0.07] | 0[0.86] | 0[0.66] | 0[0.42] | 0[0.60] |
| 182 | D | 1[0.56] | 1[0.95] | 1[0.64] | 1[0.92] | 1[0.87] | 1[0.79] |
| 183 | V | 0[0.39] | 1[0.01] | 0[1.00] | 0[0.20] | 0[0.28] | 0[0.40] |
| 184 | E | 1[0.46] | 1[0.84] | 1[0.44] | 1[0.86] | 1[0.47] | 1[0.58] |
| **185** | **Q** | **1[0.35]** | **1[0.82]** | **1[0.84]** | **1[0.90]** | **1[0.78]** | **1[0.76]** |
| 186 | I | 1[0.38] | 1[0.38] | 0[0.58] | 1[0.72] | 1[0.47] | 1[0.34] |
| 187 | K | 1[0.33] | 1[0.80] | 1[0.86] | 1[0.84] | 1[0.53] | 1[0.65] |
| 188 | Y | 0[0.31] | 1[0.19] | 0[0.66] | 1[0.58] | 0[0.14] | 1[0.01] |
| 189 | V | 0[0.29] | 1[0.04] | 0[0.46] | 0[0.20] | 0[0.41] | 0[0.36] |
| **190** | **E** | **1[0.45]** | **1[0.92]** | **1[0.24]** | **1[0.82]** | **1[0.63]** | **1[0.60]** |
| 191 | K | 1[0.35] | 1[0.95] | 1[0.68] | 1[0.84] | 1[0.41] | 1[0.59] |
| 192 | E | 1[0.16] | 1[0.81] | 1[0.44] | 1[0.84] | 1[0.42] | 1[0.54] |
| 193 | V | 0[0.48] | 1[0.09] | 0[1.00] | 0[0.66] | 0[0.49] | 0[0.64] |
| 194 | V | 1[0.31] | 1[0.89] | 1[0.20] | 1[0.92] | 1[0.31] | 1[0.50] |
| 195 | V | 0[0.49] | 1[0.45] | 0[0.84] | 1[0.04] | 0[0.29] | 0[0.26] |
| 196 | P | 0[0.10] | 1[0.87] | 1[0.04] | 1[0.76] | 1[0.46] | 1[0.46] |
| 197 | I | 1[0.10] | 1[0.40] | 1[0.02] | 1[0.78] | 1[0.40] | 1[0.41] |
| 198 | V | 0[0.02] | 0[0.03] | 0[0.44] | 1[0.10] | 0[0.40] | 0[0.24] |
| 199 | T | 1[0.28] | 1[0.38] | 1[0.80] | 1[0.80] | 1[0.36] | 1[0.53] |
| 200 | G | 1[0.89] | 1[0.83] | 1[0.60] | 1[0.92] | 1[0.83] | 1[0.77] |
| **201** | **F** | **1[0.95]** | **1[0.28]** | **0[0.20]** | **1[0.76]** | **1[0.17]** | **1[0.31]** |
| 202 | T | 1[0.99] | 1[0.42] | 1[0.28] | 1[0.76] | 1[0.35] | 1[0.46] |
| **203** | **H** | **1[0.99]** | **1[0.69]** | **1[0.48]** | **1[0.92]** | **1[0.77]** | **1[0.72]** |
| **204** | **K** | **1[1.00]** | **1[0.76]** | **1[0.04]** | **1[0.72]** | **1[0.41]** | **1[0.46]** |
| **205** | **F** | **1[0.98]** | **1[0.46]** | **0[0.82]** | **1[0.90]** | **1[0.81]** | **1[0.52]** |
| **206** | **V** | **1[0.99]** | **1[0.22]** | **0[0.24]** | **0[0.12]** | **0[0.30]** | **0[0.20]** |
| 207 | A | 1[0.97] | 1[0.61] | 1[0.24] | 1[0.72] | 1[0.39] | 1[0.47] |
| **208** | **K** | **1[0.96]** | **1[0.65]** | **1[0.28]** | **1[0.76]** | **1[0.48]** | **1[0.52]** |
| 209 | W | 1[0.59] | 1[0.50] | 1[0.30] | 1[0.80] | 1[0.31] | 1[0.45] |

| | | | | | | | |
|---|---|---|---|---|---|---|---|
| 210 | D | 1[0.96] | 1[0.84] | 1[0.70] | 1[0.86] | 1[0.58] | 1[0.67] |
| 211 | I | 0[0.63] | 0[0.48] | 0[1.00] | 0[0.60] | 0[0.80] | 0[0.79] |
| 212 | R | 1[0.93] | 1[0.85] | 1[0.46] | 1[0.82] | 1[0.63] | 1[0.64] |
| 213 | E | 1[0.93] | 1[0.85] | 0[0.20] | 1[0.88] | 1[0.70] | 1[0.59] |
| 214 | V | 1[0.94] | 1[0.42] | 0[0.82] | 1[0.26] | 0[0.10] | 0[0.06] |
| 215 | P | 1[0.91] | 1[0.68] | 1[0.82] | 1[0.88] | 1[0.72] | 1[0.74] |
| **216** | **R** | **1[0.96]** | **1[0.94]** | **1[0.70]** | **1[0.84]** | **1[0.72]** | **1[0.73]** |
| 217 | P | 1[0.97] | 1[0.54] | 1[0.24] | 1[0.78] | 1[0.26] | 1[0.44] |
| 218 | V | 1[0.99] | 1[0.06] | 0[0.46] | 0[0.26] | 0[0.58] | 0[0.40] |
| 219 | V | 1[1.00] | 1[0.58] | 1[0.42] | 1[0.78] | 1[0.32] | 1[0.49] |
| **220** | **K** | **1[1.00]** | **1[0.79]** | **1[1.00]** | **1[0.86]** | **1[0.71]** | **1[0.77]** |
| **221** | **Y** | **1[0.94]** | **1[0.40]** | **0[0.62]** | **1[0.64]** | **1[0.07]** | **1[0.17]** |
| 222 | V | 1[1.00] | 1[0.53] | 1[0.42] | 1[0.74] | 1[0.03] | 1[0.36] |
| **223** | **G** | **1[1.00]** | **1[0.57]** | **1[0.44]** | **1[0.88]** | **1[0.75]** | **1[0.69]** |
| **224** | **K** | **1[0.96]** | **1[0.89]** | **1[1.00]** | **1[0.88]** | **1[0.73]** | **1[0.78]** |
| 225 | Q | 1[0.97] | 1[0.86] | 1[1.00] | 1[0.92] | 1[0.76] | 1[0.81] |
| **226** | **E** | **1[0.89]** | **1[0.80]** | **1[0.86]** | **1[0.82]** | **1[0.71]** | **1[0.73]** |
| 227 | E | 1[0.85] | 1[0.79] | 1[0.78] | 1[0.90] | 1[0.77] | 1[0.76] |
| 228 | I | 1[0.83] | 1[0.49] | 1[0.24] | 1[0.76] | 1[0.38] | 1[0.46] |
| **229** | **E** | **1[0.71]** | **1[0.76]** | **1[1.00]** | **1[0.92]** | **1[0.61]** | **1[0.74]** |
| 230 | V | 0[0.66] | 1[0.16] | 0[0.86] | 0[0.74] | 0[0.66] | 0[0.72] |
| 231 | E | 1[0.57] | 1[0.80] | 1[0.46] | 1[0.92] | 1[0.52] | 1[0.62] |
| 232 | V | 0[0.63] | 1[0.22] | 0[1.00] | 0[0.18] | 0[0.25] | 0[0.37] |
| 233 | P | 1[0.08] | 1[0.83] | 1[0.24] | 1[0.80] | 1[0.56] | 1[0.54] |
| **234** | **Q** | **1[0.60]** | **1[0.86]** | **1[0.86]** | **1[0.88]** | **1[0.66]** | **1[0.73]** |
| 235 | V | 1[0.06] | 1[0.09] | 0[0.82] | 1[0.06] | 1[0.01] | 0[0.15] |
| **236** | **K** | **1[0.58]** | **1[0.92]** | **1[0.44]** | **1[0.88]** | **1[0.70]** | **1[0.68]** |
| 237 | F | 1[0.73] | 1[0.79] | 1[0.22] | 1[0.92] | 1[0.33] | 1[0.52] |
| 238 | V | 1[0.77] | 1[0.47] | 0[0.84] | 1[0.18] | 0[0.15] | 0[0.11] |
| **239** | **D** | **1[0.91]** | **1[0.95]** | **1[1.00]** | **1[0.92]** | **1[0.84]** | **1[0.84]** |
| **240** | **K** | **1[0.97]** | **1[0.95]** | **1[0.66]** | **1[0.90]** | **1[0.63]** | **1[0.71]** |
| 241 | V | 0[0.52] | 1[0.24] | 0[0.64] | 1[0.14] | 1[0.04] | 0[0.09] |
| 242 | V | 1[0.96] | 1[0.67] | 0[0.26] | 1[0.14] | 0[0.01] | 1[0.03] |
| **243** | **E** | **1[0.96]** | **1[0.94]** | **1[0.06]** | **1[0.92]** | **1[0.76]** | **1[0.68]** |
| 244 | H | 1[0.95] | 1[0.79] | 0[0.30] | 1[0.84] | 1[0.44] | 1[0.46] |
| **245** | **E** | **1[0.97]** | **1[0.91]** | **1[0.68]** | **1[0.90]** | **1[0.77]** | **1[0.76]** |
| 246 | V | 1[0.98] | 1[0.82] | 1[0.64] | 1[0.90] | 1[0.80] | 1[0.76] |
| 247 | V | 1[0.81] | 1[0.29] | 0[0.64] | 1[0.60] | 0[0.26] | 1[0.02] |
| 248 | V | 1[1.00] | 1[0.81] | 1[0.22] | 1[0.82] | 1[0.74] | 1[0.65] |
| **249** | **D** | **1[0.99]** | **1[0.89]** | **1[1.00]** | **1[0.90]** | **1[0.80]** | **1[0.82]** |
| 250 | T | 1[0.98] | 1[0.46] | 1[0.58] | 1[0.80] | 0[0.05] | 1[0.37] |
| 251 | I | 1[0.97] | 1[0.40] | 1[0.42] | 1[0.78] | 0[0.34] | 1[0.23] |
| **252** | **E** | **1[0.94]** | **1[0.91]** | **1[0.68]** | **1[0.88]** | **1[0.71]** | **1[0.73]** |
| 253 | K | 1[0.94] | 1[0.93] | 1[1.00] | 1[0.90] | 1[0.50] | 1[0.71] |
| **254** | **K** | **1[0.97]** | **1[0.69]** | **1[0.82]** | **1[0.92]** | **1[0.27]** | **1[0.59]** |
| 255 | V | 0[0.65] | 1[0.19] | 0[0.84] | 0[0.16] | 0[0.38] | 0[0.39] |
| 256 | P | 1[0.84] | 1[0.89] | 1[0.62] | 1[0.90] | 1[0.85] | 1[0.78] |
| 257 | K | 1[0.77] | 1[0.91] | 1[0.84] | 1[0.88] | 1[0.63] | 1[0.72] |
| 258 | I | 0[0.05] | 1[0.42] | 0[1.00] | 1[0.66] | 0[0.29] | 0[0.04] |
| 259 | I | 1[0.69] | 1[0.78] | 0[0.26] | 1[0.62] | 1[0.14] | 1[0.27] |
| **260** | **E** | **1[0.69]** | **1[0.94]** | **1[0.32]** | **1[0.92]** | **1[0.73]** | **1[0.69]** |
| 261 | V | 0[0.66] | 1[0.22] | 0[0.84] | 0[0.00] | 0[0.46] | 0[0.36] |
| 262 | P | 1[0.79] | 1[0.68] | 0[0.02] | 1[0.88] | 1[0.49] | 1[0.52] |
| 263 | K | 1[0.84] | 1[0.88] | 1[0.82] | 1[0.88] | 1[0.59] | 1[0.70] |
| **264** | **Y** | **0[0.15]** | **1[0.52]** | **0[0.04]** | **1[0.88]** | **1[0.17]** | **1[0.36]** |
| 265 | V | 0[0.23] | 1[0.19] | 0[0.46] | 1[0.34] | 0[0.25] | 0[0.09] |
| 266 | D | 1[0.80] | 1[0.86] | 1[0.82] | 1[0.92] | 1[0.61] | 1[0.72] |

| | | | | | | | |
|---|---|---|---|---|---|---|---|
| **267** | **E** | **1[0.82]** | **1[0.89]** | **1[0.70]** | **1[0.88]** | **1[0.14]** | **1[0.52]** |
| 268 | V | 1[0.24] | 1[0.67] | 1[0.02] | 1[0.86] | 1[0.64] | 1[0.56] |
| **269** | **K** | **1[0.89]** | **1[0.76]** | **1[0.48]** | **1[0.86]** | **1[0.66]** | **1[0.66]** |
| **270** | **Y** | **1[0.52]** | **1[0.63]** | **0[0.20]** | **1[0.90]** | **1[0.42]** | **1[0.47]** |
| 271 | V | 1[0.96] | 1[0.47] | 0[0.04] | 1[0.46] | 1[0.53] | 1[0.37] |
| 272 | W | 1[0.95] | 1[0.85] | 1[0.42] | 1[0.92] | 1[0.69] | 1[0.70] |
| **273** | **T** | **1[0.96]** | **1[0.75]** | **1[0.34]** | **1[0.58]** | **0[0.11]** | **1[0.26]** |
| 274 | P | 1[0.96] | 1[0.84] | 1[1.00] | 1[0.90] | 1[0.67] | 1[0.77] |
| 275 | V | 1[0.94] | 1[0.33] | 0[0.22] | 1[0.40] | 1[0.04] | 1[0.13] |
| **276** | **E** | **1[0.99]** | **1[0.85]** | **1[0.84]** | **1[0.88]** | **1[0.51]** | **1[0.68]** |
| **277** | **K** | **1[1.00]** | **1[0.85]** | **1[1.00]** | **1[0.90]** | **1[0.49]** | **1[0.70]** |
| 278 | I | 1[0.74] | 1[0.69] | 1[0.24] | 1[0.90] | 1[0.49] | 1[0.57] |
| 279 | V | 1[0.90] | 1[0.28] | 0[0.86] | 1[0.26] | 0[0.06] | 0[0.06] |
| 280 | H | 1[0.86] | 1[0.82] | 1[1.00] | 1[0.92] | 1[0.68] | 1[0.77] |
| 281 | V | 1[0.76] | 1[0.45] | 0[0.84] | 1[0.58] | 0[0.03] | 1[0.08] |
| **282** | **E** | **1[0.80]** | **1[0.75]** | **1[0.40]** | **1[0.72]** | **1[0.73]** | **1[0.62]** |
| 283 | R | 1[0.63] | 1[0.90] | 1[0.66] | 1[0.88] | 1[0.75] | 1[0.73] |
| 284 | F | 1[0.11] | 1[0.29] | 0[0.26] | 1[0.66] | 0[0.03] | 1[0.16] |
| 285 | V | 0[0.69] | 1[0.41] | 0[0.24] | 0[0.08] | 0[0.20] | 0[0.20] |
| 286 | P | 1[0.85] | 1[0.93] | 1[0.64] | 1[0.80] | 1[0.49] | 1[0.61] |
| 287 | V | 1[0.86] | 1[0.69] | 0[0.64] | 1[0.42] | 0[0.30] | 0[0.03] |
| 288 | F | 0[0.06] | 1[0.50] | 0[0.24] | 1[0.62] | 0[0.29] | 1[0.06] |
| 289 | D | 1[0.90] | 1[0.93] | 1[0.84] | 1[0.92] | 1[0.89] | 1[0.84] |
| 290 | V | 1[0.95] | 1[0.57] | 0[0.62] | 0[0.02] | 0[0.14] | 0[0.13] |
| 291 | S | 1[0.98] | 1[0.72] | 1[0.46] | 1[0.90] | 1[0.56] | 1[0.64] |
| 292 | L | 1[1.00] | 1[0.62] | 1[0.04] | 1[0.76] | 1[0.08] | 1[0.34] |
| 293 | E | 1[0.98] | 1[0.92] | 1[0.80] | 1[0.92] | 1[0.92] | 1[0.84] |
| 294 | C | 1[0.92] | 1[0.55] | 1[0.44] | 1[0.80] | 1[0.43] | 1[0.53] |
| 295 | P | 1[1.00] | 1[0.67] | 1[0.50] | 1[0.84] | 1[0.58] | 1[0.63] |
| 296 | A | 1[0.99] | 1[0.59] | 1[0.02] | 1[0.74] | 1[0.26] | 1[0.39] |
| 297 | P | 1[0.99] | 1[0.78] | 1[0.82] | 1[0.90] | 1[0.88] | 1[0.82] |
| 298 | L | 1[0.99] | 1[0.40] | 1[0.46] | 1[0.80] | 1[0.21] | 1[0.45] |
| 299 | I | 1[0.32] | 0[0.01] | 0[1.00] | 1[0.32] | 0[0.62] | 0[0.31] |
| 300 | V | 1[1.00] | 1[0.64] | 0[0.00] | 1[0.80] | 1[0.53] | 1[0.52] |
| 301 | P | 1[0.99] | 1[0.78] | 1[0.26] | 1[0.86] | 1[0.48] | 1[0.57] |
| 302 | Y | 1[0.98] | 1[0.59] | 0[0.18] | 1[0.46] | 0[0.16] | 1[0.10] |
| 303 | P | 1[0.98] | 1[0.58] | 1[0.64] | 1[0.82] | 1[0.23] | 1[0.50] |
| 304 | M | 1[0.91] | 1[0.35] | 0[0.64] | 1[0.18] | 0[0.50] | 0[0.21] |
| 305 | Q | 1[0.89] | 1[0.95] | 1[0.84] | 1[0.90] | 1[0.91] | 1[0.84] |
| 306 | A | 1[0.95] | 1[0.59] | 1[0.10] | 1[0.90] | 1[0.74] | 1[0.64] |
| 307 | V | 1[0.24] | 1[0.28] | 0[0.18] | 1[0.04] | 0[0.35] | 0[0.17] |
| 308 | K | 1[0.66] | 1[0.96] | 1[1.00] | 1[0.92] | 1[0.83] | 1[0.83] |
| 309 | E | 1[0.70] | 1[0.97] | 1[1.00] | 1[0.92] | 1[0.93] | 1[0.87] |
| 310 | M | 0[0.62] | 1[0.01] | 0[1.00] | 1[0.10] | 0[0.58] | 0[0.41] |
| 311 | P | 1[0.66] | 1[0.54] | 0[0.18] | 1[0.36] | 1[0.33] | 1[0.23] |
| 312 | A | 1[0.71] | 1[0.94] | 1[0.28] | 1[0.78] | 1[0.73] | 1[0.64] |
| 313 | V | 0[0.46] | 1[0.16] | 0[1.00] | 1[0.14] | 0[0.49] | 0[0.34] |
| 314 | M | 1[0.59] | 1[0.63] | 1[0.04] | 1[0.86] | 1[0.63] | 1[0.57] |
| 315 | V | 1[0.37] | 1[0.22] | 0[1.00] | 0[0.32] | 1[0.10] | 0[0.26] |
| 316 | R | 1[0.62] | 1[0.91] | 1[0.84] | 1[0.90] | 1[0.84] | 1[0.80] |
| 317 | K | 1[0.73] | 1[0.92] | 1[0.86] | 1[0.88] | 1[0.92] | 1[0.83] |
| 318 | E | 1[0.73] | 1[0.79] | 1[1.00] | 1[0.90] | 1[0.74] | 1[0.78] |
| 319 | V | 0[0.39] | 0[0.06] | 0[1.00] | 0[0.38] | 0[0.15] | 0[0.42] |
| 320 | P | 1[0.75] | 1[0.95] | 1[0.84] | 1[0.90] | 1[0.86] | 1[0.82] |
| 321 | E | 1[0.36] | 1[0.60] | 1[0.82] | 1[0.60] | 1[0.14] | 1[0.39] |
| 322 | A | 1[0.66] | 1[0.62] | 1[0.28] | 1[0.86] | 1[0.67] | 1[0.62] |
| 323 | A | 1[0.68] | 1[0.81] | 1[1.00] | 1[0.90] | 1[0.57] | 1[0.72] |

| | | | | | | | |
|---|---|---|---|---|---|---|---|
| 324 | I | 1[0.48] | 1[0.22] | 0[0.44] | 1[0.38] | 0[0.36] | 0[0.08] |
| 325 | T | 0[0.41] | 1[0.13] | 0[0.84] | 1[0.24] | 0[0.38] | 0[0.24] |
| 326 | E | 1[0.78] | 1[0.89] | 1[1.00] | 1[0.90] | 1[0.92] | 1[0.86] |
| 327 | Q | 1[0.76] | 1[0.79] | 1[0.60] | 1[0.64] | 1[0.01] | 1[0.36] |
| 328 | G | 1[0.73] | 1[0.79] | 1[0.24] | 1[0.78] | 1[0.36] | 1[0.48] |
| 329 | F | 1[0.56] | 1[0.25] | 0[0.42] | 0[0.06] | 1[0.13] | 0[0.06] |
| 330 | D | 1[0.83] | 1[0.82] | 1[0.28] | 1[0.90] | 1[0.78] | 1[0.70] |
| 331 | I | 0[0.07] | 1[0.24] | 0[1.00] | 0[0.04] | 0[0.53] | 0[0.41] |
| 332 | V | 1[0.85] | 1[0.82] | 1[0.82] | 1[0.90] | 1[0.78] | 1[0.78] |
| 333 | S | 1[0.78] | 1[0.46] | 1[0.58] | 1[0.86] | 1[0.73] | 1[0.68] |
| 334 | P | 1[0.67] | 1[0.75] | 1[0.58] | 1[0.64] | 1[0.34] | 1[0.47] |
| 335 | P | 1[0.60] | 1[0.87] | 1[1.00] | 1[0.84] | 1[0.61] | 1[0.71] |
| 336 | G | 1[0.37] | 1[0.22] | 1[0.24] | 1[0.32] | 1[0.52] | 1[0.32] |
| 337 | T | 1[0.42] | 1[0.43] | 1[0.06] | 1[0.52] | 0[0.14] | 1[0.14] |
| 338 | L | 0[0.52] | 0[0.19] | 0[1.00] | 1[0.10] | 0[0.62] | 0[0.44] |
| 339 | R | 1[0.36] | 1[0.55] | 1[0.42] | 0[0.20] | 0[0.56] | 0[0.23] |
| 340 | V | 1[0.28] | 0[0.22] | 1[0.14] | 1[0.18] | 0[0.36] | 0[0.11] |
| 341 | S | 1[0.49] | 1[0.26] | 1[0.42] | 1[0.58] | 1[0.29] | 1[0.36] |
| 342 | D | 1[0.58] | 1[0.85] | 1[0.66] | 1[0.88] | 1[0.73] | 1[0.72] |
| 343 | E | 1[0.66] | 1[0.49] | 1[1.00] | 1[0.62] | 1[0.26] | 1[0.48] |
| 344 | Y | 1[0.23] | 0[0.41] | 0[0.64] | 0[0.66] | 0[0.69] | 0[0.68] |
| 345 | V | 0[0.40] | 0[0.45] | 0[0.62] | 0[0.28] | 0[0.39] | 0[0.45] |
| 346 | R | 1[0.87] | 1[0.86] | 1[0.68] | 1[0.76] | 1[0.83] | 1[0.73] |
| 347 | A | 1[0.91] | 1[0.37] | 1[0.24] | 1[0.46] | 1[0.32] | 1[0.33] |
| 348 | H | 0[0.08] | 0[0.33] | 0[0.64] | 0[0.54] | 0[0.48] | 0[0.56] |
| 349 | H | 1[0.73] | 1[0.21] | 0[0.44] | 1[0.36] | 1[0.49] | 1[0.23] |
| 350 | Q | 1[0.95] | 1[0.90] | 1[1.00] | 1[0.88] | 1[0.94] | 1[0.86] |
| 351 | F | 1[0.95] | 1[0.71] | 1[0.46] | 1[0.82] | 1[0.66] | 1[0.65] |
| 352 | A | 1[0.90] | 1[0.59] | 1[0.24] | 1[0.76] | 1[0.80] | 1[0.63] |
| 353 | E | 1[0.88] | 1[0.91] | 1[1.00] | 1[0.90] | 1[0.92] | 1[0.86] |
| 354 | E | 1[0.81] | 1[0.88] | 1[1.00] | 1[0.88] | 1[0.92] | 1[0.85] |
| 355 | N | 1[0.82] | 1[0.88] | 1[0.44] | 1[0.78] | 1[0.69] | 1[0.65] |
| 356 | A | 1[0.40] | 1[0.66] | 0[0.44] | 1[0.46] | 1[0.12] | 1[0.15] |
| 357 | A | 1[0.20] | 1[0.46] | 1[0.02] | 1[0.18] | 1[0.28] | 1[0.16] |
| 358 | L | 0[0.82] | 0[0.47] | 0[0.64] | 0[0.62] | 0[0.84] | 0[0.76] |
| 359 | C | 0[0.34] | 1[0.54] | 1[0.82] | 1[0.36] | 1[0.45] | 1[0.39] |
| 360 | G | 0[0.69] | 0[0.03] | 0[0.22] | 1[0.00] | 0[0.62] | 0[0.35] |
| 361 | A | 0[0.58] | 0[0.59] | 0[0.22] | 0[0.20] | 0[0.40] | 0[0.38] |
| 362 | G | 0[0.74] | 0[0.63] | 0[0.62] | 0[0.52] | 0[0.64] | 0[0.65] |
| 363 | T | 0[0.69] | 0[0.43] | 0[0.04] | 1[0.10] | 0[0.15] | 0[0.14] |
| 364 | A | 0[0.73] | 1[0.01] | 0[0.24] | 0[0.14] | 0[0.33] | 0[0.30] |
| 365 | C | 0[0.74] | 0[0.18] | 0[0.46] | 0[0.52] | 0[0.58] | 0[0.58] |
| 366 | G | 1[0.37] | 1[0.83] | 1[0.82] | 1[0.74] | 1[0.57] | 1[0.62] |
| 367 | A | 0[0.48] | 1[0.07] | 0[0.42] | 0[0.46] | 0[0.33] | 0[0.43] |
| 368 | P | 1[0.70] | 1[0.90] | 1[0.84] | 1[0.86] | 1[0.88] | 1[0.80] |
| 369 | Q | 1[0.77] | 1[0.91] | 1[0.84] | 1[0.56] | 1[0.73] | 1[0.64] |
| 370 | K | 1[0.71] | 1[0.95] | 1[0.46] | 1[0.86] | 1[0.78] | 1[0.71] |
| 371 | E | 1[0.75] | 1[0.88] | 1[0.84] | 1[0.88] | 1[0.83] | 1[0.79] |
| 372 | P | 1[0.20] | 1[0.04] | 0[0.20] | 0[0.04] | 0[0.22] | 0[0.17] |
| 373 | E | 1[0.77] | 1[0.83] | 1[1.00] | 1[0.88] | 1[0.76] | 1[0.79] |
| 374 | I | 1[0.87] | 1[0.61] | 1[0.64] | 1[0.74] | 1[0.67] | 1[0.64] |
| 375 | E | 1[0.74] | 1[0.72] | 1[0.68] | 1[0.66] | 1[0.49] | 1[0.55] |
| 376 | F | 0[0.66] | 0[0.40] | 0[0.46] | 0[0.44] | 0[0.60] | 0[0.57] |
| 377 | R | 1[0.71] | 1[0.81] | 1[0.64] | 1[0.80] | 1[0.83] | 1[0.72] |
| 378 | T | 1[0.35] | 1[0.30] | 1[0.22] | 1[0.42] | 1[0.04] | 1[0.18] |
| 379 | A | 0[0.41] | 1[0.12] | 0[0.28] | 0[0.16] | 0[0.25] | 0[0.26] |
| 380 | N | 1[0.58] | 1[0.82] | 1[0.60] | 1[0.82] | 1[0.83] | 1[0.72] |

| | | | | | | | |
|---|---|---|---|---|---|---|---|
| 381 | Q | 1[0.49] | 1[0.79] | 1[0.50] | 1[0.64] | 1[0.79] | 1[0.62] |
| 382 | M | 0[0.24] | 1[0.01] | 0[0.82] | 1[0.06] | 0[0.40] | 0[0.31] |
| 383 | K | 1[0.56] | 1[0.82] | 1[1.00] | 1[0.80] | 1[0.78] | 1[0.76] |
| 384 | E | 1[0.45] | 1[0.87] | 1[1.00] | 1[0.88] | 1[0.88] | 1[0.82] |
| 385 | A | 0[0.35] | 0[0.12] | 0[0.48] | 0[0.18] | 0[0.10] | 0[0.26] |
| 386 | R | 1[0.42] | 1[0.82] | 1[0.44] | 1[0.50] | 1[0.57] | 1[0.48] |
| 387 | A | 0[0.11] | 1[0.26] | 1[0.18] | 1[0.34] | 1[0.22] | 1[0.19] |
| 388 | A | 0[0.28] | 1[0.19] | 0[0.46] | 1[0.24] | 0[0.09] | 0[0.07] |
| 389 | A | 0[0.34] | 1[0.03] | 0[0.10] | 0[0.10] | 0[0.02] | 0[0.13] |
| 390 | Q | 1[0.32] | 1[0.74] | 1[0.32] | 1[0.82] | 1[0.88] | 1[0.68] |
| 391 | E | 1[0.35] | 1[0.83] | 1[1.00] | 1[0.82] | 1[0.89] | 1[0.79] |
| 392 | G | 0[0.04] | 1[0.40] | 0[0.18] | 1[0.12] | 1[0.03] | 0[0.00] |
| 393 | H | 1[0.22] | 1[0.58] | 1[0.20] | 1[0.10] | 1[0.29] | 1[0.16] |
| 394 | L | 0[0.39] | 0[0.41] | 0[0.44] | 0[0.16] | 0[0.68] | 0[0.48] |
| 395 | E | 1[0.51] | 1[0.84] | 1[1.00] | 1[0.78] | 1[0.75] | 1[0.74] |
| 396 | R | 1[0.41] | 1[0.82] | 1[0.46] | 1[0.68] | 1[0.69] | 1[0.59] |
| 397 | G | 0[0.41] | 0[0.04] | 0[0.20] | 0[0.46] | 0[0.33] | 0[0.40] |
| 398 | G | 0[0.30] | 0[0.03] | 1[0.40] | 0[0.00] | 0[0.18] | 0[0.08] |
| 399 | S | 1[0.24] | 1[0.58] | 1[0.62] | 1[0.34] | 1[0.48] | 1[0.39] |
| 400 | F | 0[0.66] | 0[0.50] | 0[0.64] | 0[0.34] | 0[0.64] | 0[0.58] |
| 401 | V | 0[0.75] | 0[0.70] | 0[0.44] | 0[0.74] | 0[0.79] | 0[0.77] |
| 402 | S | 0[0.04] | 0[0.00] | 1[0.20] | 1[0.32] | 1[0.40] | 1[0.23] |
| 403 | Q | 1[0.45] | 1[0.47] | 1[0.48] | 1[0.54] | 1[0.72] | 1[0.53] |
| 404 | V | 0[0.60] | 0[0.46] | 0[0.62] | 0[0.24] | 0[0.35] | 0[0.43] |
| 405 | G | 0[0.19] | 1[0.09] | 1[0.44] | 1[0.10] | 0[0.07] | 1[0.01] |
| 406 | T | 1[0.24] | 1[0.41] | 0[0.14] | 1[0.72] | 1[0.59] | 1[0.44] |
| 407 | E | 1[0.41] | 1[0.86] | 1[1.00] | 1[0.80] | 1[0.88] | 1[0.79] |
| 408 | S | 1[0.15] | 1[0.64] | 1[0.20] | 1[0.70] | 1[0.66] | 1[0.53] |
| 409 | Q | 1[0.42] | 1[0.83] | 1[0.84] | 1[0.76] | 1[0.85] | 1[0.74] |
| 410 | V | 0[0.47] | 0[0.18] | 0[0.04] | 1[0.02] | 0[0.41] | 0[0.24] |
| 411 | P | 0[0.14] | 1[0.63] | 1[0.06] | 1[0.24] | 1[0.18] | 1[0.15] |
| 412 | R | 1[0.33] | 1[0.86] | 1[0.82] | 1[0.62] | 1[0.74] | 1[0.64] |
| 413 | P | 0[0.05] | 1[0.68] | 1[0.02] | 1[0.42] | 1[0.28] | 1[0.25] |
| 414 | A | 0[0.38] | 1[0.30] | 1[0.36] | 1[0.46] | 1[0.19] | 1[0.24] |
| 415 | A | 0[0.27] | 1[0.32] | 0[0.28] | 1[0.48] | 1[0.34] | 1[0.21] |
| 416 | T | 1[0.24] | 1[0.58] | 1[0.64] | 1[0.70] | 1[0.78] | 1[0.63] |
| 417 | D | 1[0.47] | 1[0.82] | 1[1.00] | 1[0.54] | 1[0.60] | 1[0.59] |
| 418 | E | 1[0.46] | 1[0.84] | 1[1.00] | 1[0.86] | 1[0.85] | 1[0.80] |
| 419 | Q | 1[0.49] | 1[0.81] | 1[0.50] | 1[0.86] | 1[0.87] | 1[0.73] |
| 420 | A | 0[0.27] | 1[0.10] | 0[0.30] | 1[0.24] | 0[0.06] | 0[0.04] |
| 421 | E | 1[0.54] | 1[0.91] | 1[1.00] | 1[0.80] | 1[0.86] | 1[0.79] |
| 422 | G | 0[0.00] | 1[0.59] | 1[0.64] | 1[0.32] | 1[0.34] | 1[0.32] |
| 423 | S | 0[0.07] | 1[0.60] | 1[0.42] | 1[0.74] | 1[0.56] | 1[0.52] |
| 424 | A | 0[0.41] | 1[0.36] | 1[0.36] | 1[0.32] | 1[0.14] | 1[0.17] |
| 425 | D | 1[0.51] | 1[0.89] | 1[0.28] | 1[0.62] | 1[0.70] | 1[0.56] |
| 426 | A | 0[0.30] | 1[0.38] | 1[0.12] | 1[0.38] | 1[0.34] | 1[0.24] |
| 427 | G | 0[0.11] | 1[0.46] | 1[0.34] | 1[0.20] | 1[0.17] | 1[0.16] |
| 428 | D | 1[0.56] | 1[0.80] | 1[0.82] | 1[0.34] | 1[0.71] | 1[0.53] |
| 429 | K | 1[0.65] | 1[0.90] | 1[1.00] | 1[0.76] | 1[0.85] | 1[0.78] |
| 430 | Q | 1[0.56] | 1[0.86] | 1[0.84] | 1[0.66] | 1[0.90] | 1[0.72] |
| 431 | G | 1[0.06] | 1[0.38] | 1[0.22] | 1[0.02] | 1[0.27] | 1[0.11] |
| 432 | S | 1[0.32] | 1[0.64] | 1[0.42] | 1[0.78] | 1[0.77] | 1[0.63] |
| 433 | R | 1[0.55] | 1[0.86] | 1[0.24] | 1[0.64] | 1[0.86] | 1[0.62] |
| 434 | G | 0[0.04] | 1[0.26] | 0[0.36] | 1[0.12] | 0[0.16] | 0[0.11] |
| 435 | S | 1[0.22] | 1[0.53] | 1[0.22] | 1[0.78] | 1[0.68] | 1[0.56] |
| 436 | S | 1[0.29] | 1[0.67] | 0[0.18] | 1[0.76] | 1[0.88] | 1[0.58] |
| 437 | F | 0[0.39] | 0[0.13] | 0[0.62] | 0[0.36] | 0[0.42] | 0[0.47] |

```
  438 N  1[0.41] 1[0.76] 1[0.82] 1[0.72] 1[0.88] 1[0.73]
  439 S  1[0.17] 1[0.65] 1[0.64] 1[0.76] 1[0.81] 1[0.67]
  440 E  1[0.43] 1[0.91] 1[1.00] 1[0.88] 1[0.93] 1[0.84]
  441 G  1[0.00] 1[0.49] 0[0.16] 1[0.36] 1[0.54] 1[0.28]
  442 E  1[0.87] 1[0.91] 1[1.00] 1[0.76] 1[0.92] 1[0.81]
  443 V  0[0.66] 1[0.47] 1[0.62] 1[0.78] 1[0.72] 1[0.60]
  444 H  1[1.00] 1[0.93] 1[1.00] 1[0.94] 1[0.81] 1[0.84]
```

************************************************************************

If there is no prediction printed above this asterisks line, you should go back and check your submission input. Symbols other than the 20-letter alphabet (case insensitive) are not allowed.

This is an automated email, please do not reply.
Please contact if you have questions or suggestions.
http://pipe.sc.fsu.edu
